# A natural history of AMR in *Klebsiella pneumoniae*: Global diversity, predictors, and predictions of evolutionary pathways

**DOI:** 10.1101/2025.09.20.677523

**Authors:** Olav N. L. Aga, Sabrina J. Moyo, Joel Manyahi, Upendo Kibwana, Iren H. Löhr, Nina Langeland, Bjørn Blomberg, Iain G. Johnston

## Abstract

Antimicrobial resistance (AMR) is a substantial and growing global health burden. Understanding, and predicting, its evolution in specific pathogens will help responses across scales from individual patient cases to large-scale policy. Here, we use global data on AMR features, predicted from 47k *Klebsiella pneumoniae* genomes, with hypercubic transition path sampling to infer the evolutionary pathways by which AMR features in *K. pneumoniae* (KpAMR) are acquired across 102 countries, territories and areas. We identify “globally consistent” evolutionary behaviours that hold across countries, and “globally divergent” behaviours including carbapenem and fluoroquinolone resistance that vary across countries. We show how these divergent dynamics covary both with public health superregion and drug use policy, and reveal competing evolutionary pathways within and between countries. Using newly-sequenced data across several decades from sub-Saharan Africa, we show that this inferred global roadmap of KpAMR evolution successfully predicts prospective evolutionary dynamics. Together, we hope that the ability to characterize and predict evolutionary dynamics of AMR acquisition, connected to socio-economic and drug policy predictors, will help strengthen our understanding of AMR evolution worldwide.

**Significance:** Antimicrobial resistance (AMR) occurs when microbial pathogens evolve resistance to the drugs we use to treat them. Our understanding of bacterial genomes and how they confer AMR is constantly expanding through beautiful and powerful work establishing large-scale global datasets. Here, we use emerging machine learning approaches with this genomic data to reveal the evolutionary dynamics that have generated AMR characters in a particular pathogen, *Klebsiella pneumoniae* (Kp), and how these dynamics are influenced by geography and drug use across the globe. This “natural history” of AMR in Kp makes predictions about which characters will evolve next for a given bacterium, and we validate these predictions with newly-sequenced data from clinical isolates from Africa, providing both past and prospective descriptions of AMR in Kp.

## Introduction

Antimicrobial resistance (AMR), the resistance of microbial pathogens to the medicines we use to treat them, is a major threat to global health. Antibiotic resistance, the resistance of bacteria to antibacterials, is an important subset of AMR. In 2019 and 2021, 1.27m and 1.14m deaths respectively were attributable to bacterial AMR (many more were associated with AMR), with highest rates in sub-Saharan Africa (Murray et al., 2022; Naghavi et al., 2024). The evolution of genetic and phenotypic features that cause AMR in bacterial pathogens is the focus of tremendous research interest, from basic biology through epidemiology, clinical studies and health policy.

*Klebsiella pneumoniae* (Kp) is a gram-negative bacterium and a major cause of nosocomial infections across settings. (Related species and subspecies of Kp form the KpSC (species complex); in this report we will focus on *K. pneumoniae sensu stricto*). Kp is found in multiple environmental niches and can both cause opportunistic infections in hospitalized patients and community acquired infections, including severe infections caused by hypervirulent strains (Wyres et al., 2020). Kp has been identified as an important trafficker for antibiotic resistance genes from different ecological niches into several other clinically important pathogenic bacteria (Wyres & Holt, 2018). Kp contributes significantly to the global burden of AMR. In 2021, 45 600 and 36 000 deaths were attributed to carbapenem and fluoroquinolone resistant Kp respectively. In 2021, Kp accounted for 12.7% of all deaths attributable to AMR in the population over five years old and 19.4% of AMR-attributable deaths in children under five years (Naghavi et al., 2024).

Given the substantial health burden associated with the evolution of AMR in Kp, large-scale data-driven research has explored the genomic structure of, and AMR features in, Kp populations (Argimón et al., 2021; Hetland et al., 2025; Holt et al., 2015; Munk et al., 2022; Wyres et al., 2020). Here, we consider a complementary perspective: the evolutionary pathways by which these features are acquired (Renz et al., 2024). Many related subquestions here can broadly be unified as: from a global perspective, in what orders are AMR features acquired over different instances of pathogen evolution? For example, given a newly observed isolate in the clinic with a given set of AMR features, can we predict which drug resistance(s) will evolve next (and thereby inform treatment guidelines accordingly)? And can we identify which extrinsic features – from physical environment, demography, drug policy, or others – influence the evolutionary acquisition of AMR?

These questions require approaches that can use the expanding volumes of AMR data to make and train descriptive and predictive evolutionary models. The factors shaping evolutionary pathways of AMR are increasingly studied with a combination of modelling and bioinformatic approaches (Blanquart, 2019), including connections with drug use (Olesen et al., 2018), the role of structure and heterogeneity in bacterial populations (Krieger et al., 2020), and evolutionary connections between AMR features (Lehtinen et al., 2019). However, the large sets of potentially coupled features involved in typical AMR systems are often challenging to address using traditional methods from evolutionary biology. For example, an often-studied dataset of *Mycobacterium tuberculosis* isolates contains susceptibility-resistance information for a panel of 10 different drugs (Casali et al., 2014). Comparative methods like the Mk (Markov k-state) model often struggle for more than around 7 potentially coupled characters under study (Johnston & Diaz-Uriarte, 2024). An alternative class of methods for studying the evolution of multiple characters – evolutionary accumulation modelling or EvAM – has emerged jointly from the cancer and evolution literatures (Diaz-Uriarte & Herrera-Nieto, 2022; Renz et al., 2024; Schill et al., 2024), but has traditionally considered relatively few characters in independent samples, while AMR research questions often involve many different drug resistance features in samples connected by phylogenetic relationships where presence may be due to inheritance from a common ancestor rather than independent acquisitions (Dewar et al., 2025; Maddison & FitzJohn, 2015; Renz et al., 2025).

Hypercubic transition path sampling (HyperTraPS) is an EvAM approach designed to incorporate phylogenetic information into the (Bayesian) inference of evolutionary pathways for multiple interacting characters (Aga et al., 2024; Greenbury et al., 2020; Johnston & Williams, 2016). Other EvAM approaches including TreeMHN (Luo et al., 2023), HyperHMM (Moen & Johnston, 2023), and HyperMk (Johnston & Diaz-Uriarte, 2024), also support phylogenetically coupled data, but we focus on HyperTraPS here because of its flexibility and support for relatively large numbers of features (Aga et al., 2024; Renz et al., 2024). In this article, we harness large-scale global datasets on AMR in Kp (KpAMR) with HyperTraPS to learn the structure and variability of evolutionary pathways of drug resistance in Kp across the globe, as well as the extrinsic factors shaping these pathways.

## Results

### Local and global inferred pathways of AMR evolution in Klebsiella

To learn and compare the likely pathways by which KpAMR is acquired in different countries, we obtained a dataset of 47 721 genomes from 102 different countries, territories and areas, and extracted details of the presence or absence of genes corresponding to each of 22 AMR classes, using PathogenWatch and Kleborate (Argimón et al., 2021) (Lam et al., 2021) (Methods; Fig. 1A-B). We refer to presence/absence of resistance genes for each of these classes as binary “resistance characters” (“character” referring to a particular property of a species in evolutionary biology). The set of characters we consider, following Kleborate, is given in the Fig. 1 caption; in this report we will use the shorthand defined there, grouping genes by drug resistance class (and Lahey class for β-lactamases) (Gupta et al., 2014; Lam et al., 2021; Tsang et al., 2024).

**Figure 1.**
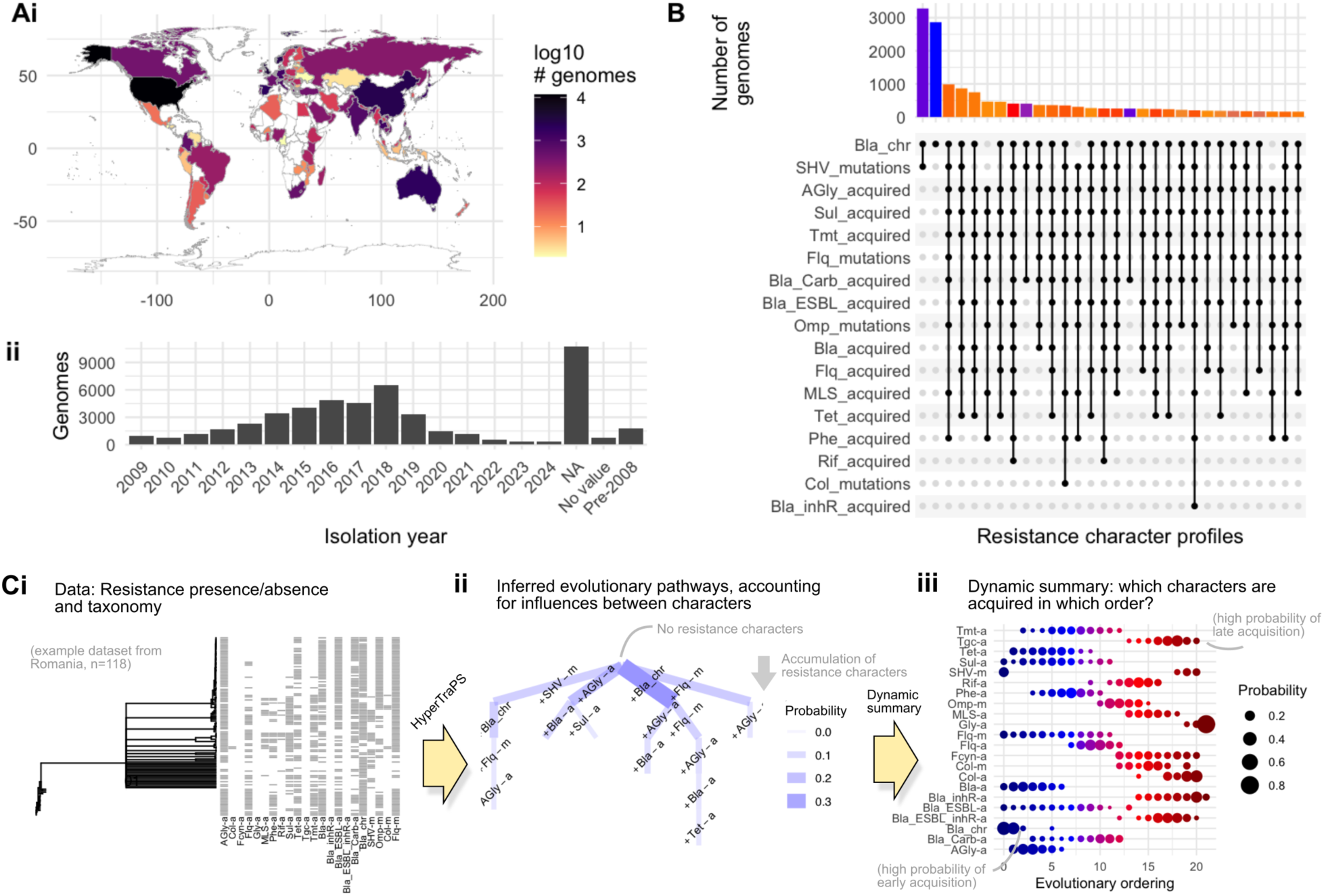
Using global genomic data on KpAMR to learn evolutionary pathways. **(A)** We use 47 721 *Klebsiella pneumoniae* (Kp) isolates from around the world, with (i) country-specific counts of isolates given by colour and years of isolation given in (ii). **(B)** UpSet plot giving counts of the 30 most common KpAMR character profiles across these isolates (bar colours give number of acquired characters). **(C) (i)** Example source data from Romania, with 118 Kp isolates connected via a putative phylogeny derived from LIN codes (see Methods) and assigned presence (dark pixel) or absence (white pixel) markers for resistance to each of 22 antibiotic groups. **(ii)** A subset of the hypercubic transition graph inferred by HyperTraPS for Romanian data. Each edge’s width gives the probability of a transition involving a single character acquisition from the upper state to the lower state. The uppermost state is the ancestral state with no AMR characters. Only early transitions above a threshold probability 0.025 are plotted for clarity. **(iii)** Summary “bubble plot” of evolutionary pathways inferred for Romanian data with HyperTraPS. The area of a bubble gives the posterior probability that a given character (row) is acquired at a given ordering in the accumulative evolutionary process. For example, *Bla_chr* is inferred to be a likely early acquisition, while *Bla_ESBL-a* likely occurs later and *Tgc_a* is inferred likely later still (or never). Colour follows a spectrum from early to late orderings. **Character names**: *AGly* (resistance to aminoglycosides); *Col* (colistin); *Fcyn* (Fosfomycin); *Flq* (Fluoroquinolones); *Gly* (Glycopeptides); *MLS* (Macrolides); *Phe* (Phenicols); *Rif* (Rifampin); *Sul* (Sulfonamides); *Tet* (Tetracyclines); *Tgc* (Tigecycline); *Tmt* (Trimethoprim); *Bla_a* (β-lactam resistance without extended spectrum or inhibitor-resistance); *Bla_inhR* (resistance to β-lactam/inhibitor combinations); *Bla_ESBL* (resistance to extended-spectrum β-lactams); *Bla_ESBL_inhR* (resistance to extended-spectrum β-lactam/inhibitor combinations); *Bla_Carb* (resistance to carbapenems); *SHV* (SHV β-lactamase with expanded enzyme activity), *Bla_chr* (SHV alleles conferring resistance to ampicillin), *Omp* (outer membrane protein). More details available via Kleborate documentation at https://github.com/klebgenomics/Kleborate. We will use *-a*: acquired (via horizontal gene transfer); *-m*: mutation; *-chr*: chromosomal (intrinsic).

For each country in our dataset, we next used hypercubic transition path sampling (HyperTraPS), a machine learning approach, to infer the “pathways” of KpAMR character acquisition: that is, the *ordered sequences* with which characters are acquired over time (see Methods). Fig. 1C illustrates this pipeline for a single country example (Romania, chosen for its moderate number of samples). HyperTraPS takes presence-absence patterns of characters on a phylogeny as input (Fig. 1Ci) and outputs an evolutionary model describing the pathways of character acquisition most supported by the data (Aga et al., 2024; Greenbury et al., 2020; Johnston & Williams, 2016; Renz et al., 2024). The full output of the inference process is a parameterised transition network describing the probability of different sequences of transitions through the state space of possible resistance characters (Fig. 1Cii). This model can be summarized, for example, as the set of posterior probabilities *P_ij_* with which character *i* is acquired at ordering *j* in a putative evolutionary process (Fig. 1Ciii). We refer to a plot where *P_ij_* is visualised with point size, as in Fig. 1Ciii, a “bubble plot”.

In Supp. Text 1, we describe some nuances involved in using HyperTraPS with these data. KpAMR characters may be acquired reversibly (for example, via plasmid gain and loss), while the model underlying HyperTraPS pictures acquisitions as irreversible. This increases the uncertainty of inference and challenges the inference of character *interactions*, but evolutionary *orderings* – which characters are likely present when another is acquired – can generally be robustly inferred (demonstrated in Supp. Fig. 1A,C and more deeply in (Johnston, 2026)). The approach does not assume an “end point” of all acquired KpAMR characters exists, that acquisitions occur one at a time, nor that populations are perfectly sampled, homogeneous, or perfectly representative of the labelled country of origin with no mixing (Diaz-Uriarte & Johnston, 2025; Krieger et al., 2020). The LINcode taxonomy linking observations (see Methods) is used to guard against potential pseudoreplication due to relatedness of samples (Boyko & Beaulieu, 2023; Dewar et al., 2025; Maddison & FitzJohn, 2015; Renz et al., 2025), but in practise (given the deep-branching nature of many relationships) has a negligible effect on inference outputs (Dauda et al., 2025; Johnston, 2026) (Supp. Fig. 1B). Finally, HyperTraPS does not assume the presence, absence, or form of interactions between features -- evolutionary dynamics can be reported for a “null model” of independent characters (with prevalence reflecting individual rates) and/or interacting characters (where rates depend on acquired characters) from the same model framework. Our focus is on characterising likely ordering dynamics, and not directly on the interactions between characters.

While Fig. 1C demonstrates our approach for a single country, Fig. 2A gives the average acquisition probabilities across all countries in our dataset, and Supp. Fig. 2 gives the corresponding summary plots for every country in the dataset. Resistance to some antibiotic groups are consistently observed across all countries: following very common presence of *Bla-chr* and *SHV-*m, resistance characters against betalactams and extended-spectrum betalactams (ESBLs) (*Bla-a*, *Bla_ESBL-a*) are often acquired earlier, while resistance to colistin (drug of last resort for multi-drug resistant *Kp*; *Col-a* and *Col-m*) is acquired later. Country-specific differences exist in the evolutionary acquisition of other characters, including resistance to carbapenems (*Bla_Carb-a)*.

**Figure 2.**
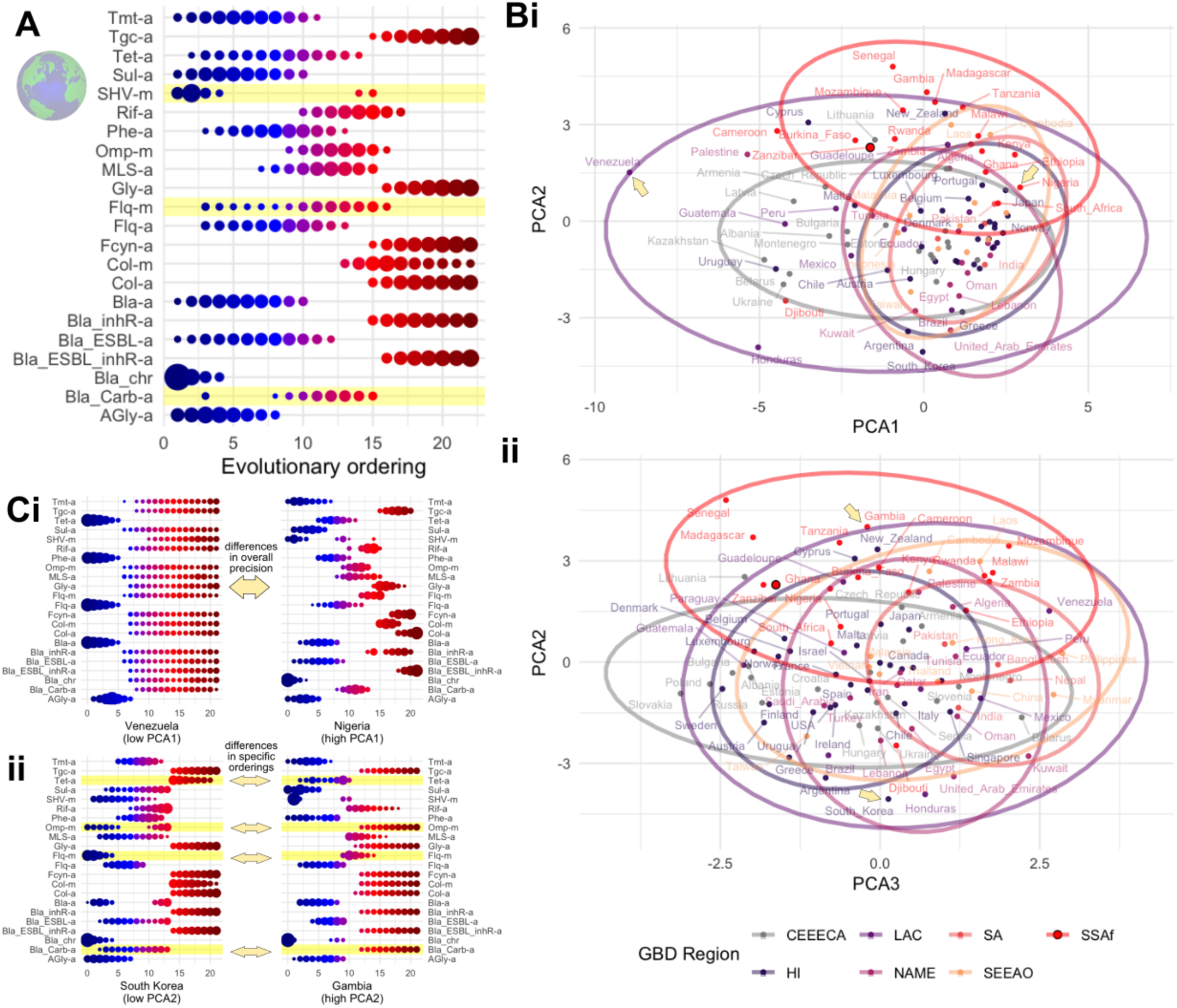
Global structure of inferred pathways of KpAMR evolution. **(A)** Global plot of summarised KpAMR evolutionary dynamics aggregated across countries. Characters displaying bimodal acquisition distributions – characteristic of distinct evolutionary pathways – are highlighted. **(B)** Principal components analysis (PCA) of country-specific “bubble plots” as in Fig. 1Ciii, reflecting inferred evolutionary dynamics of KpAMR in different countries. Individual countries are plotted on PCAs 1-2 (i) and 2-3 (ii), individually and with ellipses demarcating their corresponding GBD region (acronyms below). **(C)** Example “bubble plots” from countries (arrowed in (B)) reflecting the extremes of PCA1 (i) and PCA2 (ii). In (i), Venezuela reflects poorly characterised evolutionary dynamics, with relatively uniform probabilities everywhere; Nigeria reflects precisely characterised evolutionary dynamics, with individual characters having highly likely, precisely specified orderings. In (ii), country-specific differences in orderings emerge, including in those characters highlighted. For example, KpAMR evolution in South Korea is likely to involve *Flq-m* acquisition earlier than other characters; evolution in Gambia is likely to involve *Flq-m* acquisition later than other characters. GBD regions: CEEECA, Central Europe, Eastern Europe, Central Asia; HI, High-income; LAC, Latin American and Caribbean; NAME, North Africa and Middle East; SA, South Asia; SEEAO, South-east and East Asia and Oceania; SSAf, Sub-Saharan Africa. Zanzibar (dark outline) is from data newly sequenced in this study. Character names given in Fig. 1 caption.

In several instances, bimodality in the inferred evolutionary dynamics is observed. That is, a given character may be acquired at an earlier stage or a later stage in the evolutionary process, but not at intermediate stages. Such bimodality is characteristic of mutually repressing interactions between characters, and equivalently of several different evolutionary pathways existing in a system. Examples can readily be seen in Fig. 1C for Romania and Fig. 2A for the global mean behaviour, where *SHV-m* is likely acquired either very early or at more intermediate stages, with little probability in between. These competing dynamics are directly visible in Fig. 1Cii, where distinct, canalised pathways exist: one involving *SHV-m* as the first acquisition, one involving *Bla_chr* as the first acquisition and *SHV-m* only following rather later. Across countries (Supp. Fig. 2), bimodality is observed in several characters: some examples are in *Bla_chr* (Tanzania, Zambia); *Bla-a* (Ireland, S Korea; *Bla_ESBL-a* (Laos); and *SHV-m* (Romania and at least 15 other countries across GDB regions), reflecting multiple evolution pathways for both HGT-mediated and other KpAMR character acquisitions. Correspondingly, the country-specific inferred orderings of KpAMR characters generally correlate, but rather weakly (R^2^ = 0.41), with the simple prevalence of characters in that country’s source data (Supp. Fig. 3).

**Figure 3.**
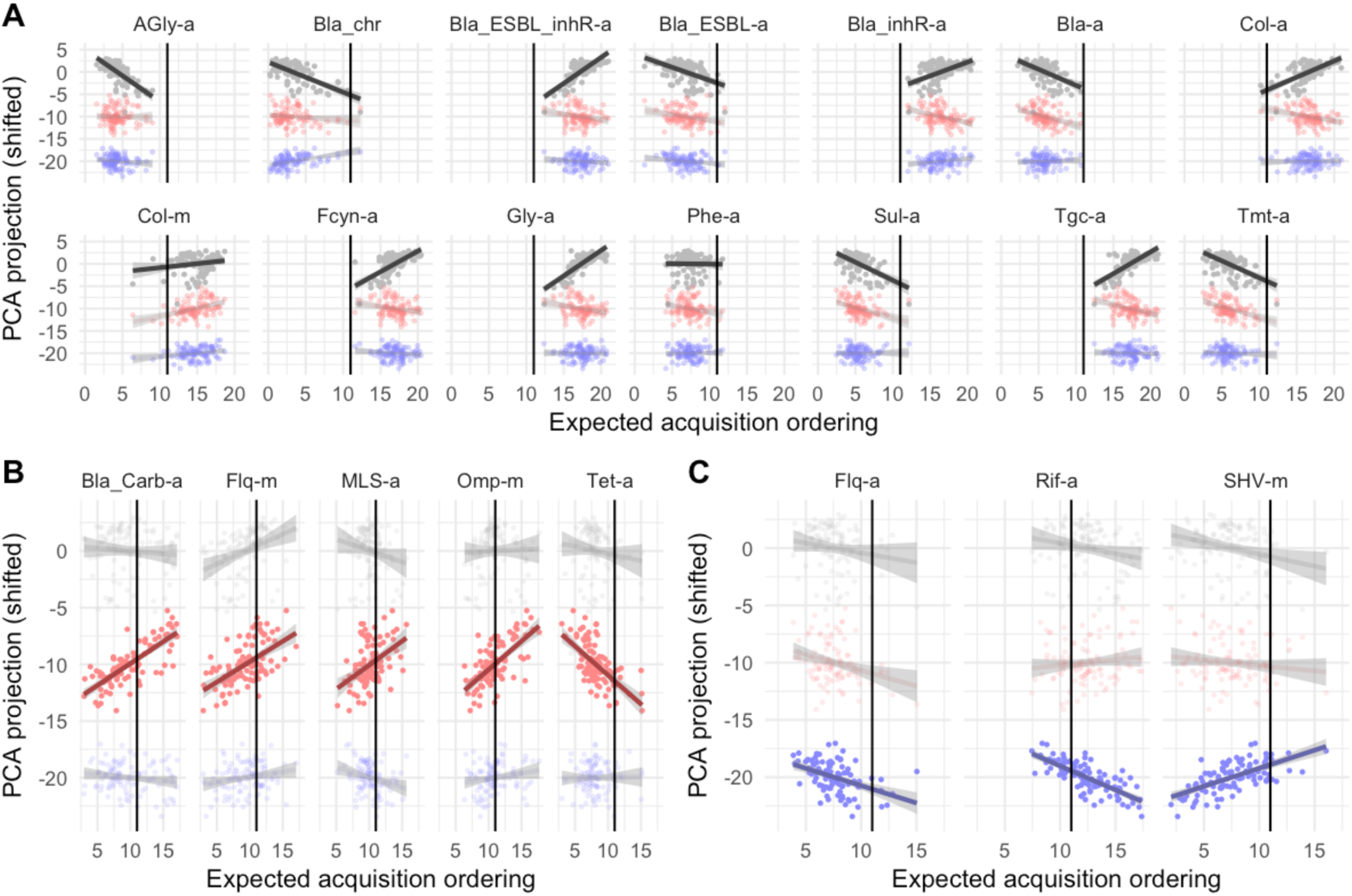
KpAMR characters determining aspects of global variability in evolutionary dynamics. Each country has an expected acquisition ordering for each KpAMR character. The plots show how these patterns of expected ordering are linked to the principal components of global KpAMR evolution variability in Fig. 2B: each point plots a country’s expected ordering for a given drug against the country’s position in PCA1 (top, grey); PCA2 (centre, red); and PCA3 (bottom, blue). **(A)** “Globally consistent” characters: those that covary tightly with PCA1. All characters here have either early or late expected orderings across all countries (either left or right of the central vertical line). Their position on PCA1 reflects the precision of the inference output. Less precise outputs (low PCA1 values, like Venezuela in Fig. 2C) will have orderings close to the central “null”; more precise outputs (high PCA1 values, like Nigeria in Fig. 2C) will have more divergent “early” and “late” orderings towards the edges of the plot. **(B-C)** “Globally divergent” characters: those that covary more tightly with PCA2 (B) or PCA3 (C). Expected orderings in these character sets span a wider range, showing sources of variability in the evolutionary dynamics that are independent of PCA1 (and hence less influenced by observational uncertainty). An alternative visualisation is given in Supp. Fig. 5. Character names given in Fig. 1 caption.

Across subsets of countries, interactions between our KpAMR resistance characters are observed (Supp. Fig. 4). For example, in around a quarter of cases, the acquisition of tigecycline resistance (*Tgc_a*) is inferred to promote the acquisition of glycopeptide resistance (*Gly_a*), and the acquisition of sulfonamide resistance (*Sul_a*) is inferred to promote the acquisition of trimethoprim resistance (*Tmt_a*). This latter observation reflects an independently-known connection: *sul* and *dfr* genes providing *Sul_a* and *Tmt_a* resistance respectively are often genetically linked, and mechanistically act on connected steps in the folate synthesis pathway (Hipólito et al., 2023; Sköld, 2001). In some cases, the acquisition of MLS resistance (*MLS_a*) is inferred to repress the acquisition of carbapenem resistance (*Bla_carb_a*), suggesting possible interactions that could be clinically exploited in combination therapies (see Discussion). However, the inference of interactions between reversibly acquired features is challenging using HyperTraPS (Johnston, 2026), so these observations must be regarded with some caution.

**Figure 4.**
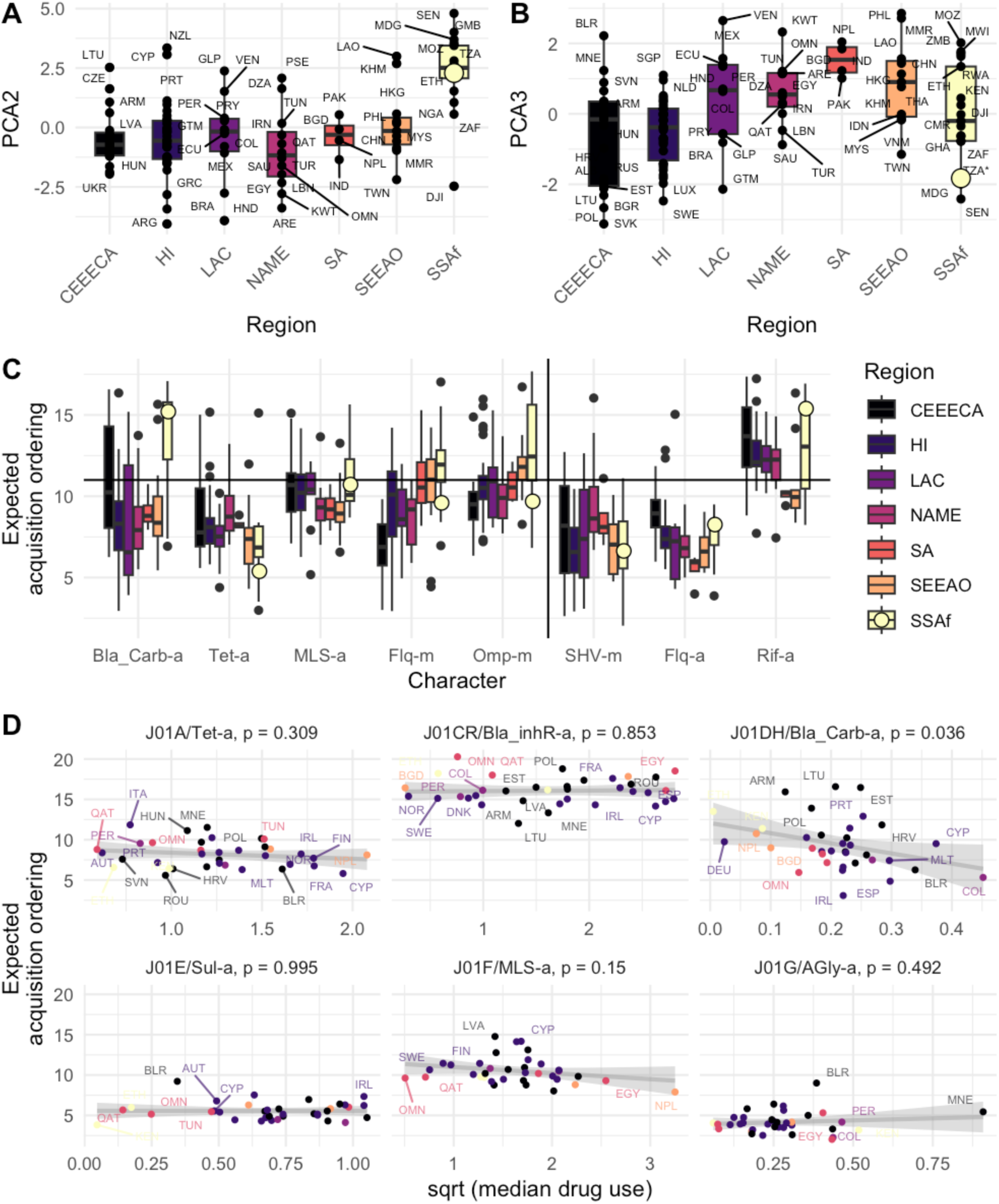
Geographical and drug policy influence on inferred AMR evolutionary pathways. **(A-B)** Distributions of country-specific PCA projection values across different GBD regions for (A) PCA2 and (B) PCA3. In (A), sub-Saharan African countries show systematically higher positions in PCA2 than other regions. **(C)** Inferred acquisition orderings of KpAMR characters by GBD region. The left five characters covary with PCA2; the right three covary with PCA3. Carbapenem resistance and OMP mutations are inferred to be acquired substantially later in sub-Saharan Africa; rifampicin and fluoroquinolone resistance are inferred to be acquired substantially earlier in S, SE, E Asia and Oceania. New Zanzibar data from this study is shown by large circles. **(D)** Connecting AMR character acquisition propensity with drug treatment regimens across countries. Median (across observed years 2016-2021) per-capita drug use in different drug classes (J codes), with expected acquisition ordering of associated resistance characters. “Globally consistent” characters *Tet-a*, *Bla_inhR-a*, *Sul-a, AGly-a* were associated with PCA1 in the global analysis and display little correlation between evolutionary ordering and drug use levels. “Globally divergent” characters *Bla_Carb-a* and *MLS-a* were associated with PCA2 in the global analysis and display stronger correlations, with earlier evolution of resistance linked to higher levels of drug use. Character names given in Fig. 1 caption; p-values are from linear regression against a null hypothesis of zero slope.

While Supp. Fig. 2 gives a detailed “roadmap” of KpAMR evolution across countries, it is hard to immediately extract global-scale insights from this representation. We therefore next considered how to compare these results in a reduced-dimensionality picture.

### Global consistency in evolutionary acquisition of resistance to a subset of drug families

To proceed, we used principal components analysis (PCA) to embed the high-dimensional set of posterior probabilities in a 3D space (Fig. 2B). Here, the axes of highest variance in the inferred evolutionary pathways are identified and used to naturally lay out the individual country outputs. Comparison of country-by-country behaviour along these axes can then reveal major sources of variability (and similarity) in inferred KpAMR evolution across countries. We underline that migration means that samples from one country likely represent a mixture of bacteria that have evolved in, and outside, that country, and that perfect separation between intrinsic dynamics of countries is therefore not possible with these data (Supp. Text 1).

Inferred evolutionary pathways for Kp across countries consistently involve early acquisition of characters conferring resistance to aminoglycosides, sulfonamides, trimethoprim, and betalactams (*Agly*, *Sul*, *Tmt* and *Bla* characters). Aminoglycosides and sulfonamides were both discovered and entered clinical use early in the mid 19^th^ century. Resistance properties inferred to be acquired at later orderings include extended spectrum beta-lactam and phenicol resistance, acquired via mobile genetic elements (*Bla_ESBL-a*, *Phe-a*). Acquisitions inferred to be acquired late across countries include inhR-driven resistance (*Bla_inhR-a*, *Bla_ESBL_inhR-a*), and resistance to later- and final-line drugs including colistin, fosfomycin, tigecyclin, and glycopeptides (*Col-a*, *Fcyn-a*, *Tgc-a*, *Gly-a*). These form a set of “globally consistent” evolutionary observations.

Country-to-country differences in evolutionary acquisition of resistance to carbapenems, fluoroquinolones, tetracycline, and others

The general patterns above appear across evolutionary dynamics across countries and regions, with the first principal component of inferred Kp variability (accounting for 23% of the overall variance) simply reporting the precision with which these patterns can be characterised (Fig. 2Ci, Fig. 3). In countries supporting less precise inference (for example, with limited datasets), the expected ordering of these characters is more uniform and closer to the average null hypothesis of all characters behaving equally; in countries with more precise output the expected orderings diverge to early and late values (Fig. 2Ci, Fig. 3).

Once variability in this precision is accounted for (by considering the first principal component), variability corresponding to other differences in the inferred dynamics can be extracted. Several characters covary comparatively little with the (precision-aligned) PCA1 and instead show stronger covariance with PCA2 or PCA3, suggesting that there is genuine country-to-country variability in these characters after controlling for differences in observational noise (Fig. 2Cii, Fig. 3, Supp. Fig. 5). PCA2 (accounting for 15% of overall variance) displays strong covariance with carbapenem, fluoroquinolone, MLS, and tetracycline resistance and OMP mutation (*Bla_Carb-a*, *Flq-m*, *MLS-a*, *Tet-a, Omp-m*,). PCA3 (accounting for 8% of overall variance) displays covariance with *Flq-a*, *Rif-a*, *SHV-m*. None of these “globally divergent” characters display the “null-to-extreme” divergence seen in those characters most closely related to PCA1, suggesting genuine country-to-country differences in evolutionary dynamics, rather than observational differences, are responsible for this variability.

**Figure 5.**
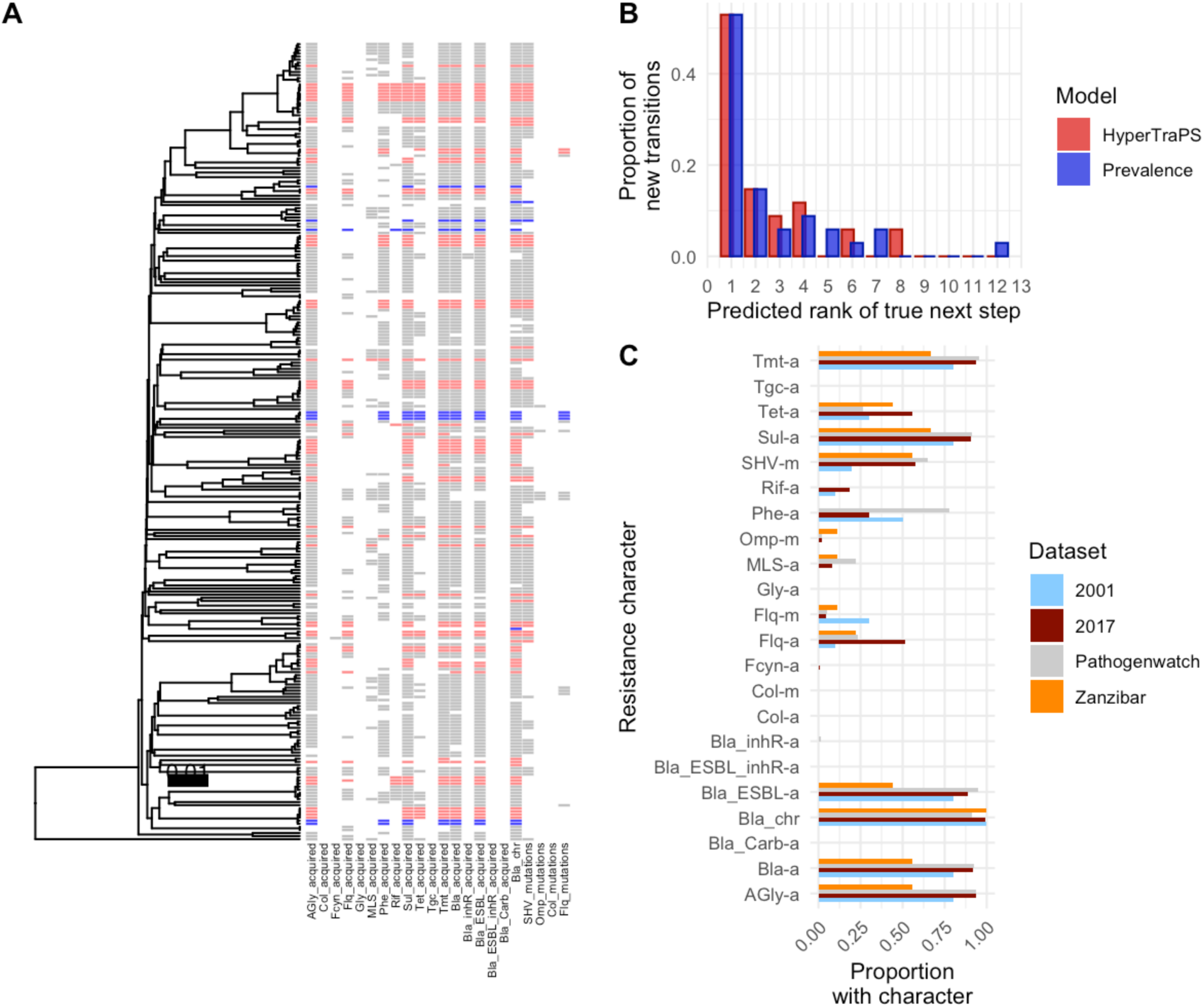
Validating prospective predictions of evolutionary transitions with newly sequenced KpAMR data. **(A)** Pathogenwatch (grey) and newly sequenced 2001-2 (blue) and 2017-8 (red) genome properties from Tanzania, connected by a phylogeny estimated from average nucleotide identity (ANI). More details in Supp. Fig. 6. **(B)** For each independent transition in the set of newly sequenced observations, we used HyperTraPS trained on existing data to predict which characters would likely evolve next given the ancestral state. The bars show the predicted rankings of the characters that were truly observed to evolve in the data. The trained model (red) usually predicted top rankings for the characters that were truly next acquired, marginally outperforming a model based purely on character prevalence (blue). **(C)** Proportion of Tanzanian isolates in Kleborate and new datasets (uncorrected for relatedness) where different KpAMR characters are present. Character names given in Fig. 1 caption.

### Geographical correlates with Kp evolutionary dynamics

To explore the sources of this evolutionary variability in more detail, we asked whether geographical region was linked to different variants of the inferred evolutionary dynamics (after accounting for observational differences). We first asked whether countries from different regions showed systematic differences in their PCA2 and PCA3 projections – corresponding to different orderings of the “globally divergent” country-specific characters above (Fig. 4). We found that sub-Saharan Africa was a dramatic outlier in PCA2, involving notably later acquisition of carbapenem and fluoroquinolone resistance and OMP mutation than the other regions (Fig. 4A, C). This ordering result agrees with the continuous-time history of treatment in the region, where carbapenem and fluoroquinolone use was established later (especially for children) than in other regions. Other regions were broadly consistent in PCA2 but showed substantial differences in PCA3. These differences correspond mainly to acquired resistance to fluoroquinolone and rifampicin, with substantially earlier inferred acquisition in South-Southeast-East Asia and Oceania, and substantially later in Central and Eastern Europe, Central Asia, and sub-Saharan Africa (Fig. 4B, C).

### Policy predictors of variance in AMR evolutionary dynamics

We next asked whether known policy of antibiotic use influence the inferred evolutionary dynamics from our analysis. We gathered statistics on drug use from 2016-2021 in countries available from the WHO GLASS surveillance programme (*Global Antimicrobial Resistance and Use Surveillance System (GLASS) Report 2022*, 2022) and looked at correlations between per-capita drug use statistics and the inferred evolutionary dynamics of resistance to those drugs (Fig. 4D). We found that for those drug types associated with PCA1 (where variability in inferred dynamics is largely determined by observation uncertainty) there was little correlation between drug use and the ordering of resistance acquisition. However, for carbapenems, associated with PCA2 (more non-observational country-to-country variability), there was a link between drug usage statistics and inferred resistance evolution, with carbepenam resistance (*Bla-carb-a*) more generally present before other acquisitions in countries with high drug use rates. This observation could arise through increased *Bla-carb-a* acquisition rate, or (perhaps more likely) a reduced loss rate of the *Bla-carb-a* feature once acquired (see Supp. Text 1). The direction of this correlation was also observed in MLS, also associated with PCA2, though this relationship did not show statistical support at the p < 0.05 level. This pattern of correlations is consistent with the picture above: acquisition of some (PCA1) KpAMR characters is relatively constant across countries and circumstances, while other (PCA2) characters are more plastic and depend more tightly on country- and region-specific drivers. We note, however, that the GLASS data only cover a subset of years for which bacterial samples exist in our data (Fig. 1A) and that KpAMR evolution is not constrained only within these dats (Supp. Text 1), so our drug use measure is only an imperfect estimate of the strength of any selective pressure shaping the dynamics.

### Testing predictions of AMR evolutionary dynamics with newly sequenced, cross-decadal data

One motivating application of evolutionary inference to AMR problems is the ability to predict the next steps in evolutionary pathways (Renz et al., 2024). A question of potentially direct clinical relevance is: given a clinically observed isolate with a given set of AMR characters, which character(s) will likely evolve next – and can treatment guidelines be adapted accordingly? Previous studies of AMR with HyperTraPS have demonstrated its ability to predict properties of evolutionary transitions in a withheld subset of the overall dataset, following the common training-test machine learning paradigm (Aga et al., 2024). Here, we are in the unique position of using independently obtained and Kp genome data to test the predictions of the evolutionary models trained above.

We sequenced Kp isolates from patients with bloodstream infections in 2001-2 and 2017-8 in Tanzania and 2015-6 in Zanzibar, collected in previous studies (Blomberg et al., 2007; Moyo et al., 2020; Onken et al., 2015, 2024) (Fig. 5; see Methods). Bioinformatic analysis placed these new genomes in a phylogenetic tree with a subset of existing genomes from the training data, including their AMR profiles (Fig. 5A, Supp. Fig. 6). Using only transitions independent of those in the training data (see Methods), we tested the predictions of the HyperTraPS model trained on data from Tanzania. Specifically, for each new transition, we queried the trained model about likely next steps from the ancestral state, and recorded the predicted ranks of the next steps actually present in the transition (Fig. 5B). By far the most common outcome was that the true next step was ranked first (predicted to be most likely). For these data, the trained model outperformed a naïve model based only on the relative prevalence of KpAMR traits only in a subset (18%) of cases – this is because the newly-observed data showed little evidence of higher-order effects (including bimodal ordering distributions and interactions between characters), which HyperTraPS can readily capture (demonstrated in previous applications to AMR data (Aga et al., 2024; Renz et al., 2025) but a prevalence-only model cannot (hence the variability in Supp. Fig. 3).

This analysis demonstrates that within-country predictions of future evolutionary dynamics are possible given a trained HyperTraPS model. We also asked whether the model could predict evolutionary dynamics at the regional scale. We use Zanzibar as a test case here – a region of Tanzania that was not explicitly represented in the original Kleborate training dataset. Our findings would predict KpAMR in Zanzibar to fall into the sub-Saharan Africa set of behaviours, with high values on PCA2 and associated signatures of late carbapenem resistance and others (Fig. 4A). To this end, we applied HyperTraPS inference to newly sequenced isolates from Zanzibar. The resulting inferred dynamics clearly place Zanzibar within the set of sub-Saharan Africa behaviours (Fig. 2B), with high values on PCA2 arising from later acquisition of carbapenem resistance, and high values on PCA3 arising from later acquisition of rifampicin resistance (Fig. 4A-B).

We also explored the representation of different KpAMR characters in the snapshot genome sequences alone, without connecting with evolutionary dynamics. Fig. 5C shows the relative prevalence of acquired KpAMR characters in existing and newly-sequenced data, not accounting for phylogenetic relatedness (and without representation of an explicit non-AMR control group). With this sampling, most characters showed an increase from 2001-2 to 2015-6, particularly in mobile genetic acquisition of fluoroquinolone resistance, while the relative prevalence of acquired phenicol resistance and fluoroquinolone resistance mutations decreased. Proportions of these characters were in broad agreement with known coarser-grained observations from Tanzania (Mapunjo et al., 2025; Sangeda et al., 2021) and existing Tanzanian genomes from Pathogenwatch, but we observed more acquired rifampicin resistance and lower acquired phenicol resistance than in existing samples. Our samples from Zanzibar showed lower relative proportions of aminoglycoside and β-lactam resistance, and higher prevalence of outer membrane protein mutations.

We have embedded the trained HyperTraPS KpAMR models in an online Shiny app https://stochasticbiology.shinyapps.io/amr-predict/. Here, a country of interest can be specified, and the KpAMR profile of a sample given by specifying presence or absence of each of our characters. The app will then give the inferred probabilities of each next possible character acquisition, as well as a transition graph describing the likely subsequent pathways for up to five acquisitions in the future (Supp. Fig. 7).

## Discussion

Powerful data-driven approaches to study the evolution of AMR are emerging in parallel with large genomic datasets (Argimón et al., 2021; Hetland et al., 2025; Holt et al., 2015; Munk et al., 2022; Wyres et al., 2020). Most of these studies focus on the current genomic structure of pathogen populations. In this research we have attempted to take a complementary perspective, using large-scale genome data while focusing on the (inferred) evolutionary dynamics that produce these genomic patterns. We believe that this new way of working has substantial power to support other data-driven approaches, particularly in the prediction of unseen and future evolutionary behaviours of pathogens (Renz et al., 2025).

The core technology we use, HyperTraPS, is an instance of evolutionary accumulation modelling or EvAM (Diaz-Uriarte & Herrera-Nieto, 2022; Diaz-Uriarte & Johnston, 2024). EvAM has previously been used to learn AMR pathways, including in HIV (Beerenwinkel et al., 2005) and multidrug resistance in tuberculosis (Aga et al., 2024; Greenbury et al., 2020; Moen & Johnston, 2023; Renz et al., 2024). One question that has remained is to what extent such results reflect universal, general behaviors, versus a response to the specific selective pressures arising from a given country’s drug use (for example). Our global comparison suggests a form of answer to this question. A subset of the KpAMR characters we consider appear to behave similarly across countries regardless of specific selective pressure, drug regimes, and other differences (those aligned with PCA1). Another set of characters (those aligned with PCA2-3) evolves in a way that is more dependent on country- and region-specific influences, which we have shown include public health superregions and drug use levels – although certainly other factors, from population change to the prevalence of other diseases, will also influence these dynamics (Blanquart, 2019; Olesen et al., 2018). While these results are specific to *Kp*, and the characters we consider, this combination of global and country-specific behaviors may reasonably hold across other pathogens.

Inferring evolutionary dynamics of multiple, coupled traits across groups is a challenging problem. There will be countless, multiscale selective influences that impact the evolution of specific lineages of *Kp* around the globe. Our perspective attempts to “coarse-grain” away from these (unobservable) specific pressures and consider their aggregated effect on large-scale genomic properties in populations. This perspective is not without difficulties. In Supp. Text 1 we describe in more depth several resolutions to challenges including reversibility of KpAMR acquisitions (for example, via plasmid gain and loss), phylogenetic relatedness, mixing between countries, imperfect observations, and presence or absence of interactions between characters. As described there and studied in (Johnston, 2026), evolutionary orderings from HyperTraPS inference are rather robust to these issues in real data, which tend to increase uncertainty rather than systematically bias the outputs of inference. The “controlling” for uncertainty via PCA1 in our analysis is therefore expected to ameliorate the effects of these issues. As shown in Supp. Fig. 1C, the inference of *which characters are likely present when another is acquired* is congruent across our method and an alternative that, while limited in scale, can directly allow for reversibility (Johnston & Diaz-Uriarte, 2024).

In learning estimates for evolutionary dynamics, HyperTraPS attempts to infer any potential interactions between characters -- for example, demonstrating an independently known positive connection between sulfonamide and trimethoprim resistance (Hipólito et al., 2023; Sköld, 2001). Negative interactions, where the acquisition of one character makes acquiring another character less likely, could (if truly present) form the basis of a combination therapy XY, where resistance to drug X makes resistance to drug Y less likely. These interactions correspond to the phenomenon of *collateral sensitivity* in AMR, where resistance to one drug induces sensitivity to another (Pál et al., 2015; Szybalski & Bryson, 1952): for example, in *Escherichia coli*, beta-lactamase expression is linked to heightened sensitivity to colistin (among other) (Herencias et al., 2024). While not observed consistently throughout all countries in our dataset, a subset of countries showed statistical support for some such “repressive interactions” (Supp. Fig. 4). These include, for example, MLS resistance repressing the acquisition of carbapenem resistance. However, this is one aspect of inference which *is* challenged when a reversible process is modelled irreversibly (Johnston, 2026), and such indications can only suggest more detailed followup investigation to verify and explore mechanisms behind these possibly exploitable collateral interactions.

In conclusion, we hope to show that evolutionary accumulation modelling – inferring the historic evolutionary pathways of AMR acquisition, in addition to considering contemporary properties – can complement and enhance established ways of working in AMR genomic analysis. By considering the dynamics by which global KpAMR patterns have become established, a collection of research directions are opened or expanded, including the similarities and differences between countries, links with social and environmental covariates, and predictions of future dynamics. We anticipate that these methods may readily be applied to provide insight into AMR evolution in other pathogens in future.

## Methods

### Existing Klebsiella data acquisition

We obtained all 47 721 *Klebsiella pneumoniae sensu stricto* records from pathogenwatch as of March 2024 (Argimón et al., 2021). The most common isolation years were 2015-2020, with a peak at 2018 (Fig. 1A). We used AMR features (genes and mutations) reported by drug class as reported by Kleborate version 2.3 (Lam et al., 2021). Kleborate groups genes or mutations known to confer resistance to clinically relevant antibiotic groups and related phenotypes. Betalactamases are further grouped by enzyme activity (Lam et al., 2021). We consider a set of *L* = 22 drug classes for each genome, dichotomizing Kleborate output by presence of any gene or mutation related to each group. That is, if there is any gene or mutation present for a resistance phenotype, the isolate is considered resistant, otherwise it is considered susceptible. This assumption holds for most resistance phenotypes considered in this study. However, fluoroquinolone resistance (*Flq-m*) often require multiple mutations to cause resistance. Thus, the presence of these characters cannot be interpreted as a resistance phenotype, but rather an increased MIC (minimum inhibitory concentration). The isolates were divided by country based on metadata, and countries were grouped into superregions as defined by the Global Burden of Disease regional classification system (Rudd et al., 2020). A coarse-grained phylogeny was generated via LIN-codes (Life Identification Numbers: a systematic, stable quantitative system for genomic classification based on nucleotide identities (Marakeby et al., 2014), pre-computed for our dataset (Hennart et al., 2022)) using lincoding.py supplied by (Hennart et al., 2022). 2 102 genomes were excluded due to missing metadata. Any country with less than two genomes available were excluded due to the inability to construct a tree. The resistance profiles and phylogenetic tree were combined into a phylogenetic tree annotated with resistance to drug classes.

### Evolutionary pathway inference with hypercubic transition path sampling (HyperTraPS)

HyperTraPS (Aga et al., 2024) models an “evolutionary space” containing every possible state of a system involving *L* binary characters, then infers the probabilities of (Markovian) transitions between states in this space (here, involving the ordered accumulation of different AMR characters) that is most compatible with observed data (Greenbury et al., 2020; Johnston & Williams, 2016). A recent overview of the method in an AMR context is given in (Renz et al., 2024). The source data is a collection of length-*L* “barcodes” labelling presence or absence of our characters -- resistance to drug classes as defined in Kleborate -- on the tips of a phylogeny. Ancestral state reconstruction (here, assuming that AMR resistance character accumulation is rare and irreversible, so that an ancestor possesses a character if and only if all descendants possess it) is by default used to infer ancestral states, and thereby construct a set of transitions from the data. This process is used solely to guard against pseudoreplication, and for these (largely deeply-branching) datasets, the inclusion of phylogenetic detail (and hence the details of ancestral state reconstruction) have a negligible influence on the inference outputs (Supp. Fig. 1B). The resulting transitions are the input data for the HyperTraPS algorithm. The target of inference is an *L* x *L* matrix θ, where θ_*ii*_ is the base rate for acquiring character *i*, and θ_*s*j_ is the influence that acquisition of character *j* has on the base rate of acquisition of character *i* (as in mutual hazard networks (Schill et al., 2020)). Specifically, the probability of acquiring character *i* from the state corresponding to binary vector *s* is *P*(*s* gains *i*) = exp(θ_*ii*_ + ∑ _*sj*_ θ_*sj*_)/ ∑_*k*_ exp(θ_*kj*_ + ∑ *s_j_* θ*_kj_*), where ∑_*k*_ is over all characters absent in *s* (Aga et al., 2024). In this way, positive and negative interactions between the acquisition of different AMR characters is supported, so that (for example) acquisition of resistance to drug X can make acquisition of resistance to drug Y more (or less) likely. The output of the inference process is, more broadly, a transition matrix λ, with elements λ_*a→b*_describing the probability with which an isolate in state *a* will next transition to state *b*. For each of the 102 countries, territories and areas we ran HyperTraPS on three different random seeds to ensure that the results were consistent across runs (Supp. Fig. 2). Simulations were run for 10^6^ steps with 10^3^ random walkers for sampling, employing a penalised likelihood involving the HyperTraPS-estimated likelihood for a given set of transitions given a parameterisation θ, computed with the probability term above calculated over sampled paths, and a unit penalty for each non-zero parameter (see (Aga et al., 2024)). The US genomes were sub-sampled to a subset of 1000 genomes; three different subsamples clustered closer together than with any other countries in the PCA plot.

### Synthetic control study for inference of reversible characters

To determine the effect of our assumption of irreversible accumulation of AMR characters (given that the loss of plasmids is not uncommon in Kp evolution (Holt et al., 2015)), we constructed a synthetic control study. A random phylogeny over 128 tips was constructed using a birth-death process with birth rate 1 and death rate 0.5 (chosen empirically to roughly match the phylogenies from our case studies) using *phangorn* (Schliep, 2011). A synthetic accumulation process reflecting a single evolutionary pathway was simulated on this phylogeny. Different loss processes – including uniform rate over a subset of features, heterogeneous, and random rates – were simulated in different instances (details in Supp. Fig. 1A caption). The same HyperTraPS pipeline (assuming irreversibility), including ancestral state reconstruction and pathway inference, was run on both the pre-loss and post-loss datasets, and the outputs compared (Supp. Fig. 1A). We also applied HyperMk (Johnston & Diaz-Uriarte, 2024), an EvAM approach supporting reversible dynamics, to a reduced subset of the KpAMR data (HyperMk is limited in the scale of data it can analyse), showing both a tight agreement with the HyperTraPS output and that model selection criteria favour the irreversible model (Supp. Fig. 1C). These are a small-scale illustrative examples; the inference of reversible dynamics under an irreversible model is the focus of study in (Johnston, 2026).

### Global structure in inferred pathways

The outputs of HyperTraPS can be summarized as a matrix *P*, where element *P_ij_* gives the posterior probability that character *i* is acquired at ordering *j* in a putative evolutionary process from an ancestor with no AMR characters towards a final state with all AMR characters. For example, *P_32_* gives the probability that the third character is the second to be acquired in such an evolutionary process. We obtained this matrix for each country in our dataset, then performed principal components analysis (PCA) on the set of matrices (Williams et al., 2013). We observed the structure of matrices with “bubble plots”, where an array of points is plotted with areas proportional to *P_ij_*.

### Drug usage data

We collected antibiotic consumption data from 57 countries, territories and areas enrolled in the WHO Glass surveillance program in the period 2016 to 2021 (*Global Antimicrobial Resistance and Use Surveillance System (GLASS) Report 2022*, 2022). Overlaying the consumption data with the countries that had 3 or more available genomes reduced the number to 53. We use the median consumption statistics over the available time points for each country.

### New Klebsiella data acquisition

The 79 newly sequenced Kp isolates in this article are derived from a collection of studies performed in Tanzania and Zanzibar over the last three decades. These comprise bacterial isolates from (a) blood samples from pediatric patients with bloodstream infections in Dar-es-Salaam, Tanzania in 2001-2002; (b) blood samples from adult patients with bloodstream infections in Dar-es-Salaam, Tanzania in 2017-2018; (c) fecal samples from the patients in (b); and (d) blood samples from adult and pediatric patients with bloodstream infections in Mnazi Mmoja hospital, Zanzibar in 2015-2016. These cultures were sequenced by MicrobesNG (MicrobesNG, Birmingham, UK) using Illumina HiSeq technology. Assembled contigs were provided by the sequencing service, which formed the initial step for the bioinformatics pipeline below.

### New Klebsiella data analysis

The analysis pipeline begins with the contigs from the initial sequencing and assembly process. We first used *dnadiff*, which wraps *nucmer* from the MUMmer 3.0 package (Kurtz et al., 2004), to scaffold the contigs from a given isolate on the *Klebsiella* reference genome GCA_000240185.2_ASM24018v2 (Liu et al., 2012) and report statistics of the alignment. We retained only isolates with <20% unaligned bases as instances of *Klebsiella.* Combining the newly sequenced isolates with the existing Tanzanian genomes from Pathogenwatch, we then ran pairwise *dnadiff* across the combined set. We extracted the average identity across the aligned regions for each pair and recorded this as the ANI (average nucleotide identity). We then used *d* = 1-ANI as a distance measure for phylogenetic tree estimation using the unweighted pair group method with arithmetic mean (UPGMA) (Sokal & Michener, 1958) implemented in the *phangorn* (Schliep, 2011) package in R (R Core Team, 2022). We confirmed that an alternative method, neighbourhood joining (Saitou & Nei, 1987), did not give qualitatively different results for the predictions of AMR evolution. We used AMR profiles of existing and new isolates to reconstruct ancestral states across this complete tree, then identified the set of subtrees that had only newly-sequenced isolates at their tips. As transitions within such subtrees are independent of transitions in the (Pathogenwatch) training data set, we used this set of transitions as our independent test set.

### Testing predicted dynamics from the trained model on new data

We queried the model fitted from the training data to predict the likely next states from each precursor state in this independent set of transitions (Aga et al., 2024; Renz et al., 2024). We ranked the possible next steps from each state from most likely to least likely under the fitted model. We then recorded the ranks corresponding to the actual step(s) corresponding to the observed transition, and compared these to a null model where each possible transition could occur with probability given by its relative prevalence in the training data. For the Shiny app in Supp. Fig. 7, we report the probabilities of each possible next acquisition step given a provided current state, and construct the transition network starting from that current state and involving up to five further acqusitions.

## Data and code availability

The code base for this study is publically available at https://github.com/StochasticBiology/kp-evolution-inference. In addition to the software referenced above, we use R (R Core Team, 2022) with libraries *hypertrapsct* for evolutionary inference (Aga et al., 2024), *dplyr* (Wickham et al., 2023), *tidyverse* (Wickham et al., 2019), *countrycode* (Arel-Bundock et al., 2018) for data manipulation, *lme4* (Bates et al., 2015) for statistics, *phytools* (Revell, 2012) for phylogenetic analysis, and *ggplot2* (Wickham, 2016), *ggpubr* (Kassambara, 2020), *ggbeeswarm* (Clarke et al., 2016), *ggrepel* (Slowikowski, 2021), *ggupset* (Ahlmann-Eltze, 2025) for visualisation. We also use a Python script for LINcoding by Melanie Hennart, which uses *numpy* (Harris et al., 2020). **Authors’ note:** we are in the process of uploading the new Kp sequences to a repository; a link will be provided when this process is complete. In the meantime the raw FASTA files (which are the input for the pipeline) can be downloaded from https://osf.io/36r45.

## Acknowledgements

This project has received funding from the European Research Council (ERC) under the European Union’s Horizon 2020 Research and Innovation Programme [Grant agreement No. 805046 (EvoConBiO) to I.G.J.]. This project was supported by the Trond Mohn Foundation [project HyperEvol under grant agreement No. TMS2021TMT09], through the Centre for Antimicrobial Resistance in Western Norway (CAMRIA) [TMS2020TMT11], This project has received support from ERA-Net: JPI-AMR STRESST project (NFR333432). The authors are grateful to the CAMRIA collaboration, the Stochastic Biology Group at UiB, and delegates to the Nordic AMR conferences 2024 and 2026 for useful discussions.

## Supplementary Information

### Supplementary Text 1. Nuances and resolutions in applying HyperTraPS to KpAMR data

The model underlying HyperTraPS pictures acquisitions as occurring irreversibly, while KpAMR characters – particularly those acquired through plasmid exchange – may be gained and lost many times by a lineage (Renz et al., 2025). However, while reversibility increases the uncertainty of the inference process and challenges the inference of the detailed interactions between features, inferred evolutionary dynamics and particularly acquisition orderings is robust to this challenge – illustrated with some simple examples in Supp. Fig. 1A, comparison with an EvAM approach allowing for reversibility (Johnston C Diaz-Uriarte, 2024) in Supp. Fig. 1C, and the focus of study in (Johnston, 2026). This holds both for comparable loss rates across features and for distinct loss propensities, in which case the approach functions to infer the gain/loss rate ratio – which itself determines evolutionary orderings and prevalence patterns (Johnston, 2026). Generally, the outputs of HyperTraPS infer and report *which features are likely present when a given feature is gained* – a quantity that remains well-defined across irreversible and reversible cases. The congruence of this inference across reversible and irreversible approaches can be seen, for example, in Supp. Fig. 1C.

We also note that the model does not assume that a final “end point” with all characters present is ever reached (or indeed possible), instead leaving uncertain all acquisitions that are not observed in real data. HyperTraPS invokes a “emission” picture, comparable to a hidden Markov model, to connect with observations (Johnston C Williams, 2016), and therefore does not require complete information or homogeneous populations – it can naturally account for limited sampling of a broader, potentially heterogeneous population (Diaz-Uriarte C Johnston, 2025).

HyperTraPS does not *a priori* assume that the inferred evolutionary dynamics are a consequence of independent character accumulation (where each character’s observed prevalence is determined by a single acquisition rate) or interactions between characters (where the acquisition of a character is influenced by the presence of others) (Johnston, 2026). The “null model” of independence is challenged, for example, by instances of bimodality in inferred orderings – where a character may be acquired before many others or after many others, but not at intermediate stages. But the inferred evolutionary trajectories may be reported and analysed regardless of which generative model is most supported. (We note, in parallel to this point, that simultaneous acquisition of multiple features is very compatible with our approach; the resulting output simply places equal probabilities over all compatible ordering pathways.)

Our samples are connected by a phylogeny, which we estimate with a taxonomy constructed from LINcodes (see Methods). Accounting for this phylogeny allows us to control against pseudoreplication, where individuals inheriting characters from a common ancestor are interpreted as independent acquisition instances of those characters (Boyko C Beaulieu, 2023; Dewar et al., 2025; Maddison C FitzJohn, 2015; Renz et al., 2025). Our approach uses the *relative* evolutionary timings from the source data phylogeny (for example, Fig. 1Ci) to report the *orderings* of events: which acquisitions precede or follow which others, given that characters may influence each other, with the timescale of these events (see Discussion). In our case, much of the taxonomy corresponds to LINcode elements reflecting a deep timescale, with only separations within the final clusters corresponding to separation timescales under 1000 years (following a back-of-the-envelope calculation in footnote below). For many KpAMR features, a picture of independent acquisitions is then not unreasonable, as many KpAMR acquisitions may have occurred on a shorter timescale (that of modern human medicine). However, our phylogenetically-embedded approach only biases the inference in the case of strong connection between acquisition patterns and sampling (Johnston, 2026), otherwise simply increasing the uncertainty of the inferences (Dauda et al., 2025; Johnston, 2026). In many real-world EvAM cases, whether or not phylogeny is included has an effect on uncertainty but no detectable impact on estimated orderings – demonstrated for two countries in our dataset in Supp. Fig. 1B. As we control for the different uncertainties across countries in the followup PCA analysis, we use this approach to correct for potential instances of pseudoreplication.

The large majority of genomes with timing information were isolated in the period 2015-2020 (Fig. 1A). Our approach here does not directly consider the specific, real-world timings of the evolutionary transitions involved. The precise timescales of the evolutionary process are in principle addressable using a version of our approach called HyperTraPS-CT (continuous time) (Aga et al., 2024), but this requires a consistent estimation of divergence times between all isolates in the dataset. Importantly, our approach here does not completely neglect a timescale – it is implicit in the phylogenetic relationships included in the source data, and this inclusion both accounts for the temporal ordering of events and “pseudoreplication” as above. However, this approach based on relative evolutionary timings means that we cannot directly tie differences in inferred evolutionary behaviour to known real-world historical events shaping AMR profiles: shifts in drug policy, for example, or outbreaks or other events changing the effective size of the evolving population. While our results, focusing on the ordering of acquisition events, already demonstrate descriptive and predictive power in KpAMR evolution across countries, a continuous time picture on a smaller scale in the future (perhaps a single country or small set) would enable finer-grained linking between model prediction and the real-world details of policy and disease history.

Our samples are labelled by country of collection, which we treat as a covariate in our analysis. However, of course, bacteria and their hosts are not physically limited to remain within a given country. The phylogeography of Kp has been studied in depth, for example in (Han et al., 2019; Hetland et al., 2025; Wu et al., 2023) and certainly involves mixing of strains across borders. Our analysis and interpretation does not assume that observations from a country all and exclusively reflect evolution in that country alone; our regional and correlative downstream analysis accepts that mixing can readily take place and attempts to identify variability that can be observed despite this mixing (which, in the limiting case, would lead to homogeneous inferred dynamics worldwide).

-----

The terminal clusters in our LIN-derived taxonomy reflect ANI >99.84%, giving a per-site substitution proportion of at most 0.0016. Estimates for Kp substitution rates are around 1.5×10^-6^ substitutions per site per year (Aguilar-Bultet et al., 2023). A substitution proportion of 0.0016 might then be expected over a timescale of 0.0016/1.5×10^-6^ ≈ 10^3^ years, reflecting an estimate for the longest timescale of separation within the terminal clusters.

**Supplementary Figure 1.**
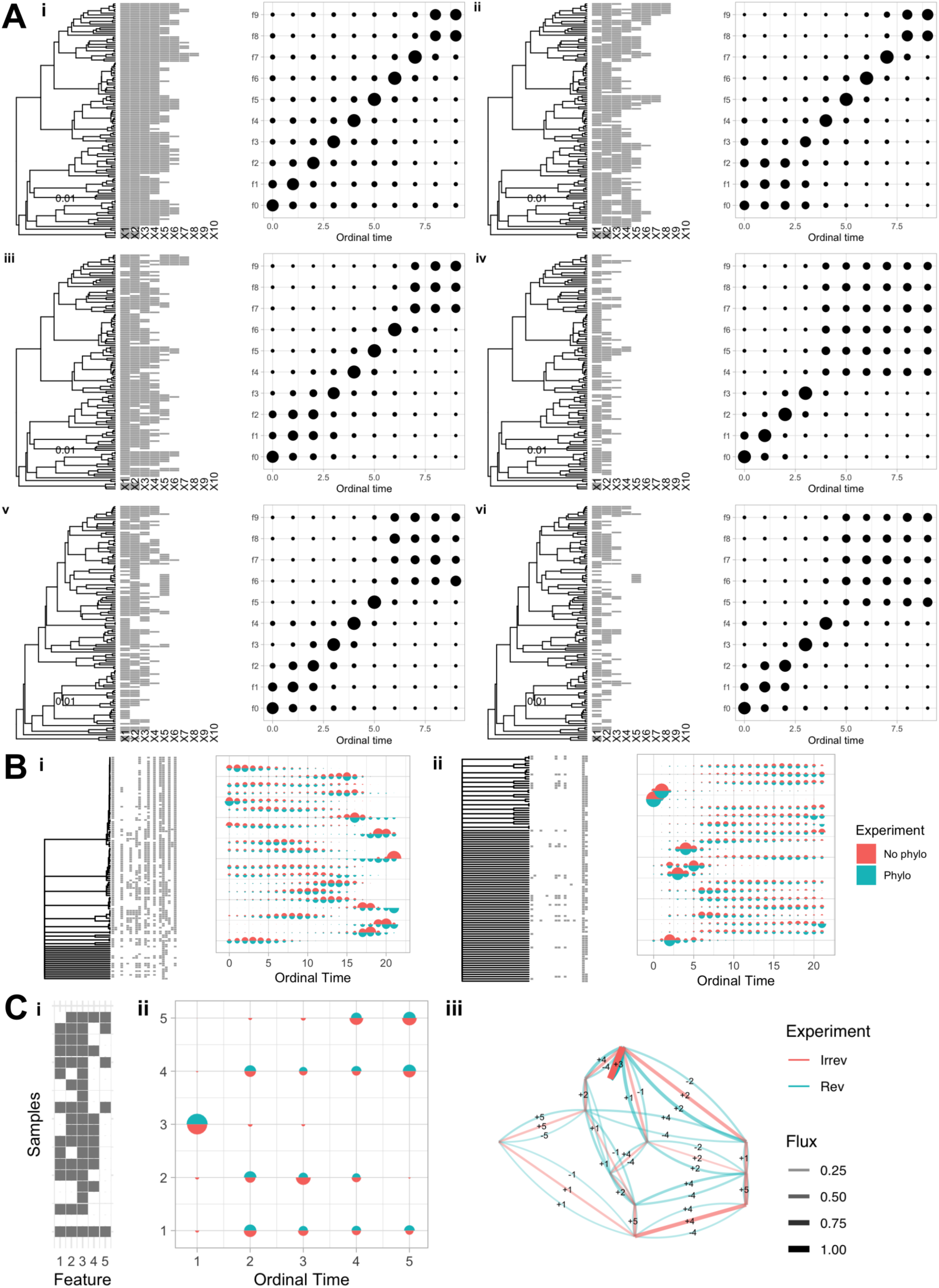
Limited issues with modelling reversible dynamics. Plots involve a dataset (left) and “bubble plot” summary of inference as in the text (right). **(A) HyperTraPS recovers accumulation orderings from reversible processes.** Synthetic datasets involving the progressive accumulation, and loss, of characters X1 to X10, with different accumulation and loss rates (i)-(vi). For (i) accumulation rate = 1 and no loss take place. For (ii)-(vi) accumulation rate = 2 and (ii) loss rate is 0.1 for features 1-5 and 0 otherwise; (iii) loss rate increases linearly from 0.05 for feature 1 to 0.5 for feature 10; (iv) loss rate increases linearly from 0.2 for feature 1 to 2 for feature 10; (v) loss rate for each feature is drawn from U(0, 0.5); (vi) loss rate for each feature is drawn from U(0, 1). The output inferred from HyperTraPS accurately reports the accumulation ordering of characters, with differences in uncertainty, and ambiguity about characters which are not observed across all samples. This small-scale case study reflects a specific instance of the picture explored more generally in (Johnston, 2026). In all cases, inferred ordering can be interpreted as reporting which features are present when another is gained (see Supplementary Text). **(B) Limited impact of neglecting phylogenies.** Datasets from (i) Romania; (ii) Senegal. Inferences shown ignoring phylogeny (cross-sectional picture) vs including phylogenetic information for ancestral state reconstruction and considering transitions down lineages. Here (and generally), many samples are related only via deep branches and are effectively independent; more recent relatedness within clades has a marginal effect on the uncertainty of posteriors but no noticeable effect on inferred orderings, in agreement with (Dauda et al., 2025; Johnston, 2026). **(C) Agreement of irreversible and reversible inference.** (i) A reduced subset of the full KpAMR dataset, involving 5 features chosen to reflect a spread of prevalences: *Bla_ESBL_a* (1), *Sul_a* (2), *Bla_chr* (3), *Tet_a* (4), *Rif_a* (5). With this reduced set, the HyperMk approach (Johnston C Diaz-Uriarte, 2024) can be used to perform inference allowing for reversibility. Both the bubble plot summary (ii) and the inferred transition network (iii) for HyperMk (“Rev”) are very similar to the HyperTraPS outputs assuming an irreversible picture (“Irrev”), with HyperMk supporting reversible versions of all the same transitions identified (also observed in (Johnston, 2026)); the Akaike Information Criterion favours the irreversible picture (393 vs 553).

**Supplementary Figure 2.**
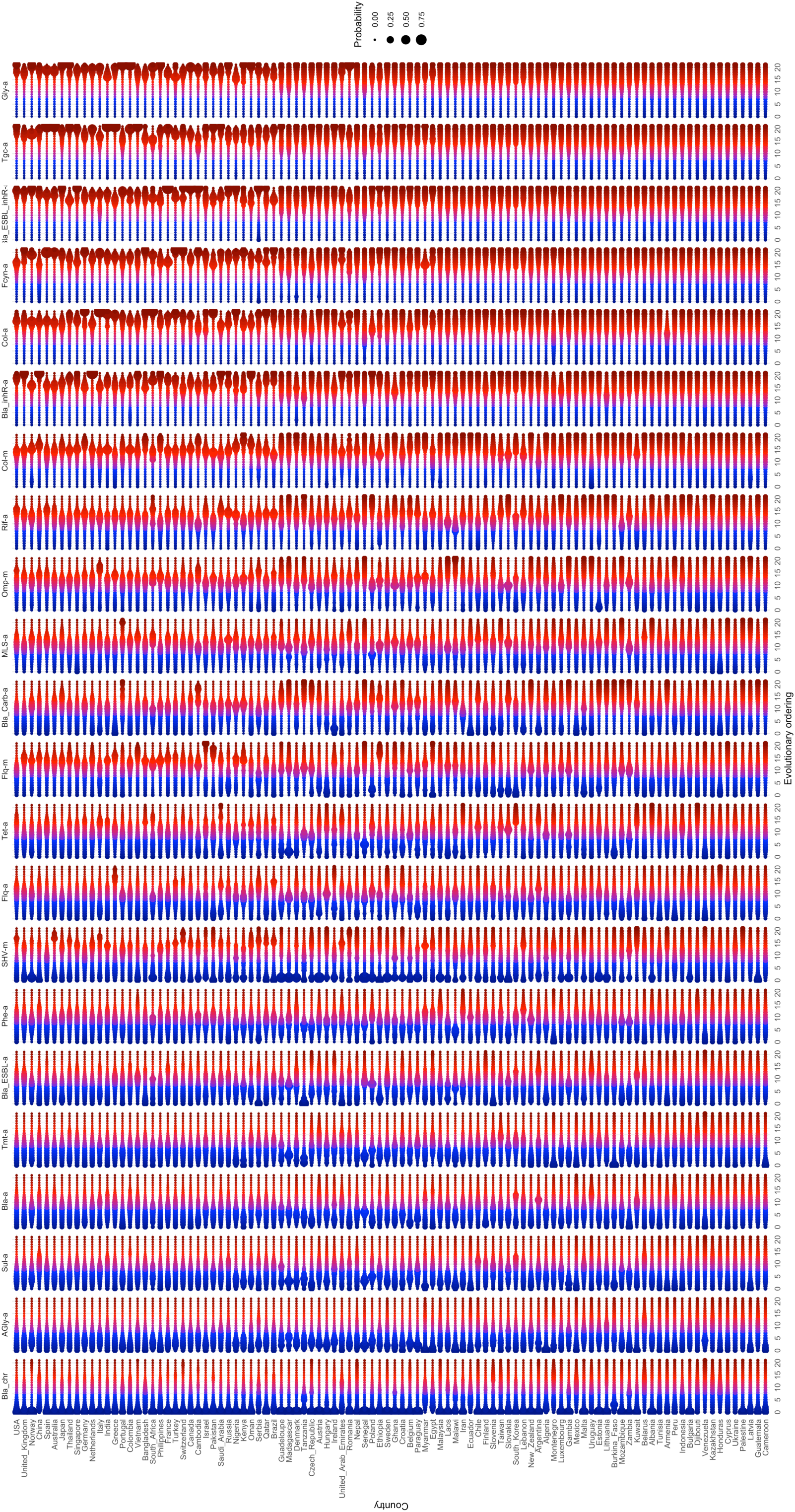
All posterior orderings of AMR character acquisitions. This “global roadmap” summarises “bubble plots” describing inferred evolutionary dynamics of KpAMR characters in different countries. Each row is a country, ordered vertically by descending number of Kp samples. Each column is a KpAMR character; the horizontal axis within columns gives evolutionary orderings (from early to late). The size of a point gives the probability that that character is acquired at that ordering in that country. For example, across the vast majority of datasets (including the top row, USA, for example), *Bla_chr* has a high probability of early acquisition and *Gly_acquired* has a high probability of late acquisition. In Cameroon, all characters have similar inferred evolutionary orderings, reflecting the limited data available to make more precise statements. Character names given in Fig. 1 caption.

**Supplementary Figure 3.**
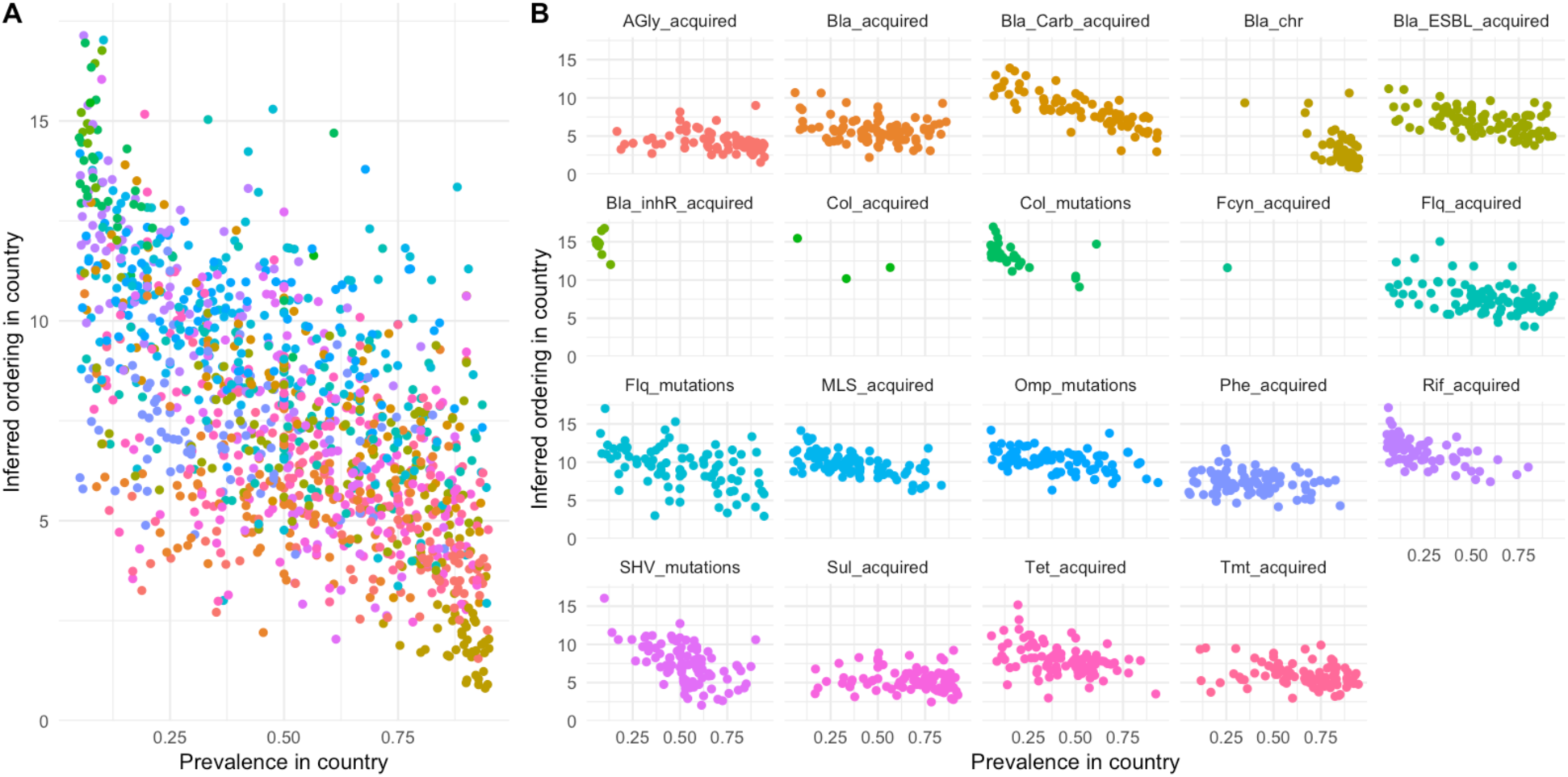
Mean posterior orderings and country-specific prevalences of AMR character acquisitions. **(A)** All KpAMR characters with a prevalence >5% and <95% together (R^2^ = 0.41). (B) Individual KpAMR characters.

**Supplementary Figure 4.**
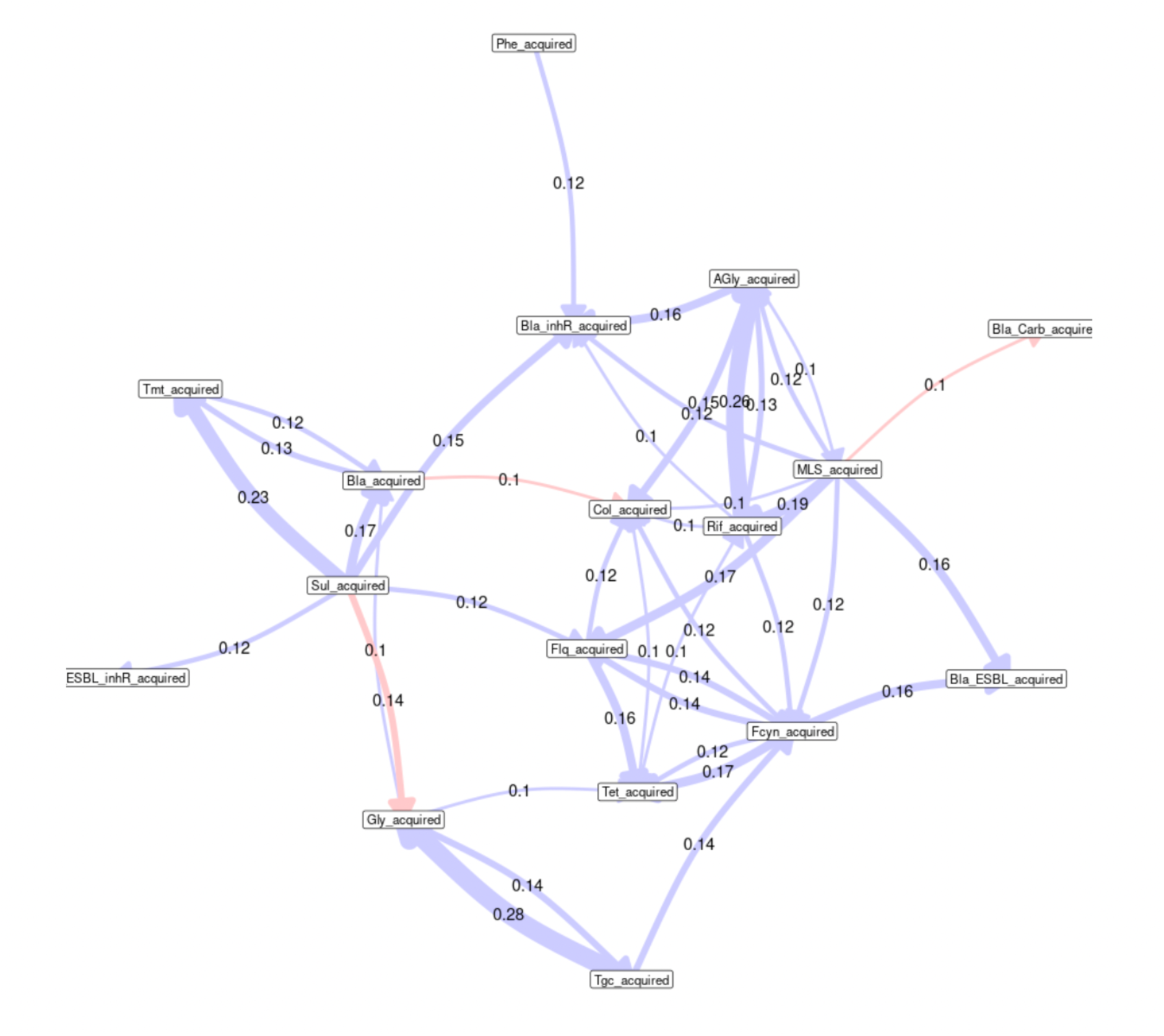
Inferred interactions between KpAMR characters. An arrow from character X to character Y reports an inferred interaction between the evolutionary acquisition of those characters. Blue arrow: acquiring X makes acquiring Y more likely (“promoting”). Red arrow: acquiring X makes acquiring Y less likely (“repressing”). The number on each arrow gives the proportion of countries in which that interaction was inferred. Character names given in Fig. 1 caption. However, as the inference of character interactions with HyperTraPS is challenging for reversible systems (Johnston, 2026), these results should be treated with caution.

**Supplementary Figure 5.**
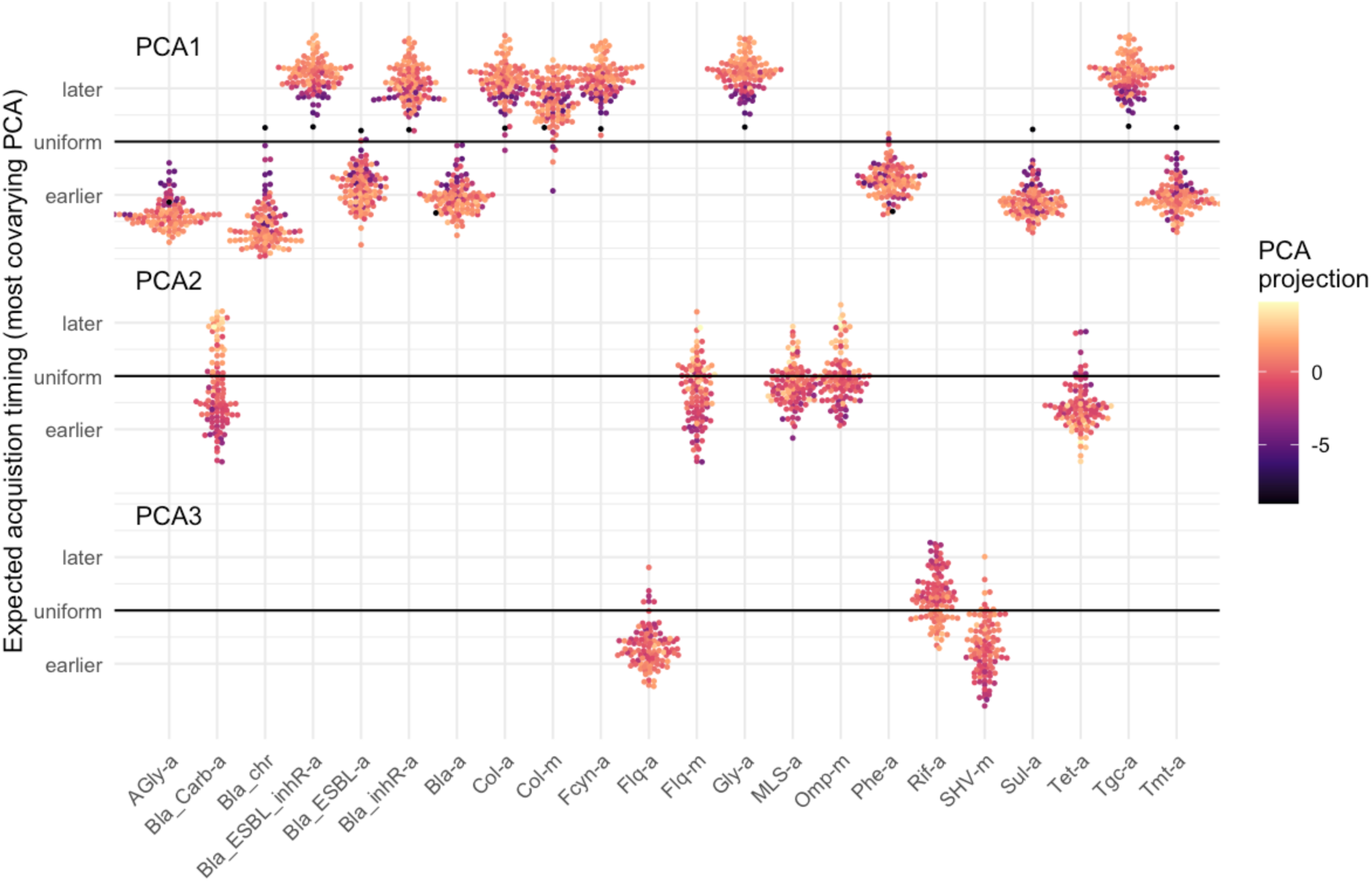
KpAMR characters determining aspects of global variability in evolutionary dynamics (alternative visualisation). Connected to Fig. 3. KpAMR characters (horizontal axis) are plotted by their expected acquisition ordering (vertical axis) for each country (points). The country’s projection on PCA1, PCA2, and PCA3, for those characters that most strongly covary with each PCA axis, are given by the colour of a point. As in Fig. 3, those characters that covary with PCA1 form a consistent spectrum linked to precision: low values of PCA1 correspond to less precise, more uniform timing, while higher values lead to more precise earlier or later timings. Characters covarying with the other PCAs display wider ranges connected to PCA1-independent variability. Character names given in Fig. 1 caption.

**Supplementary Figure 6.**
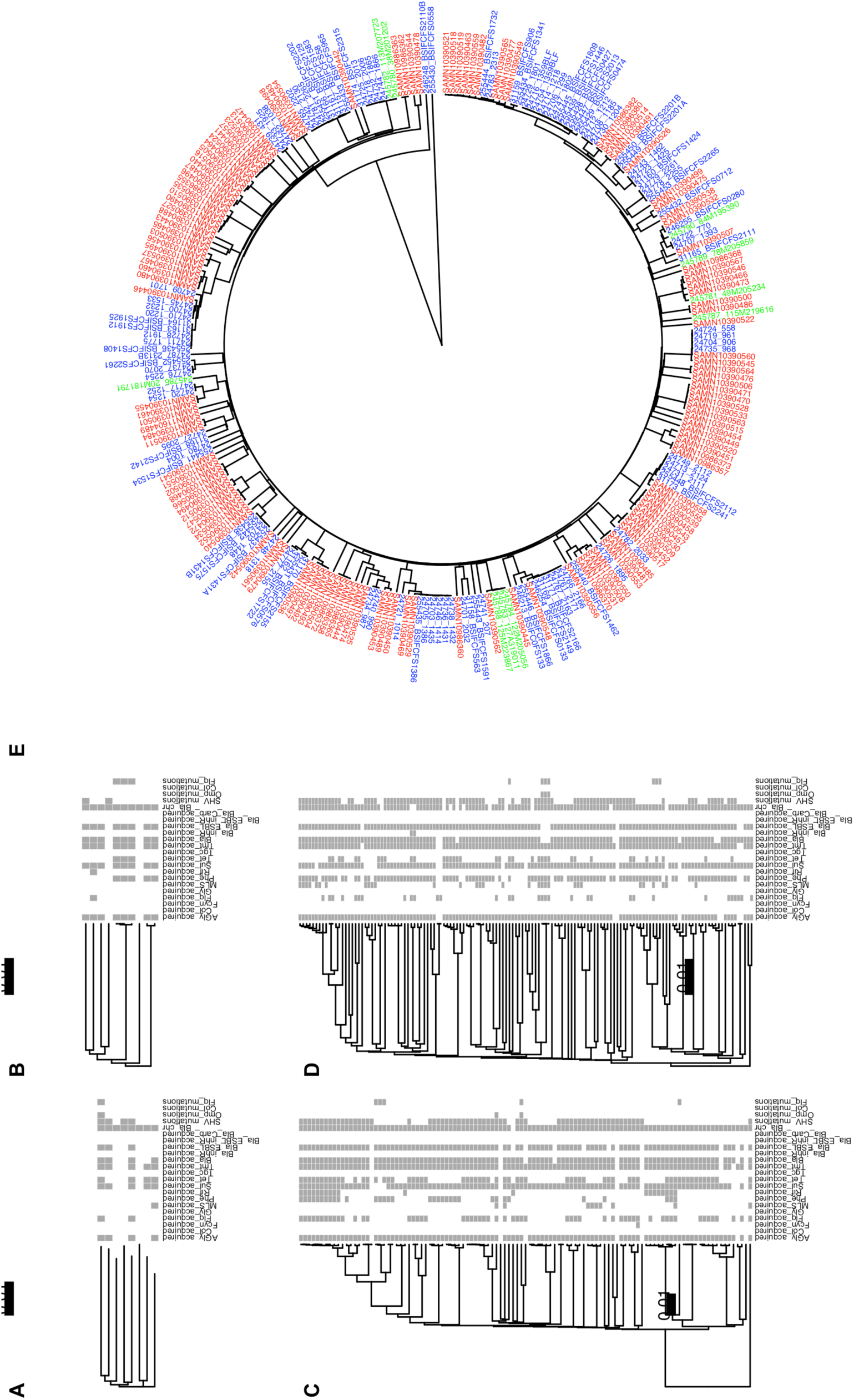
New KpAMR data from Zanzibar and Dar es Salaam, Tanzania. Phylogenies and KpAMR profiles from new datasets: **(A)** Zanzibar 2015-6. **(B)** Tanzania 2001-2. **(C)** Tanzania 2017-8. **(D)** Pathogenwatch Tanzania. **(E)** Circular projection of the phylogeny of Tanzanian isolates in Fig. 5A. Colours: green, 2001-2; blue, 2017-8; red, existing Pathogenwatch data.

**Supplementary Figure 7.**
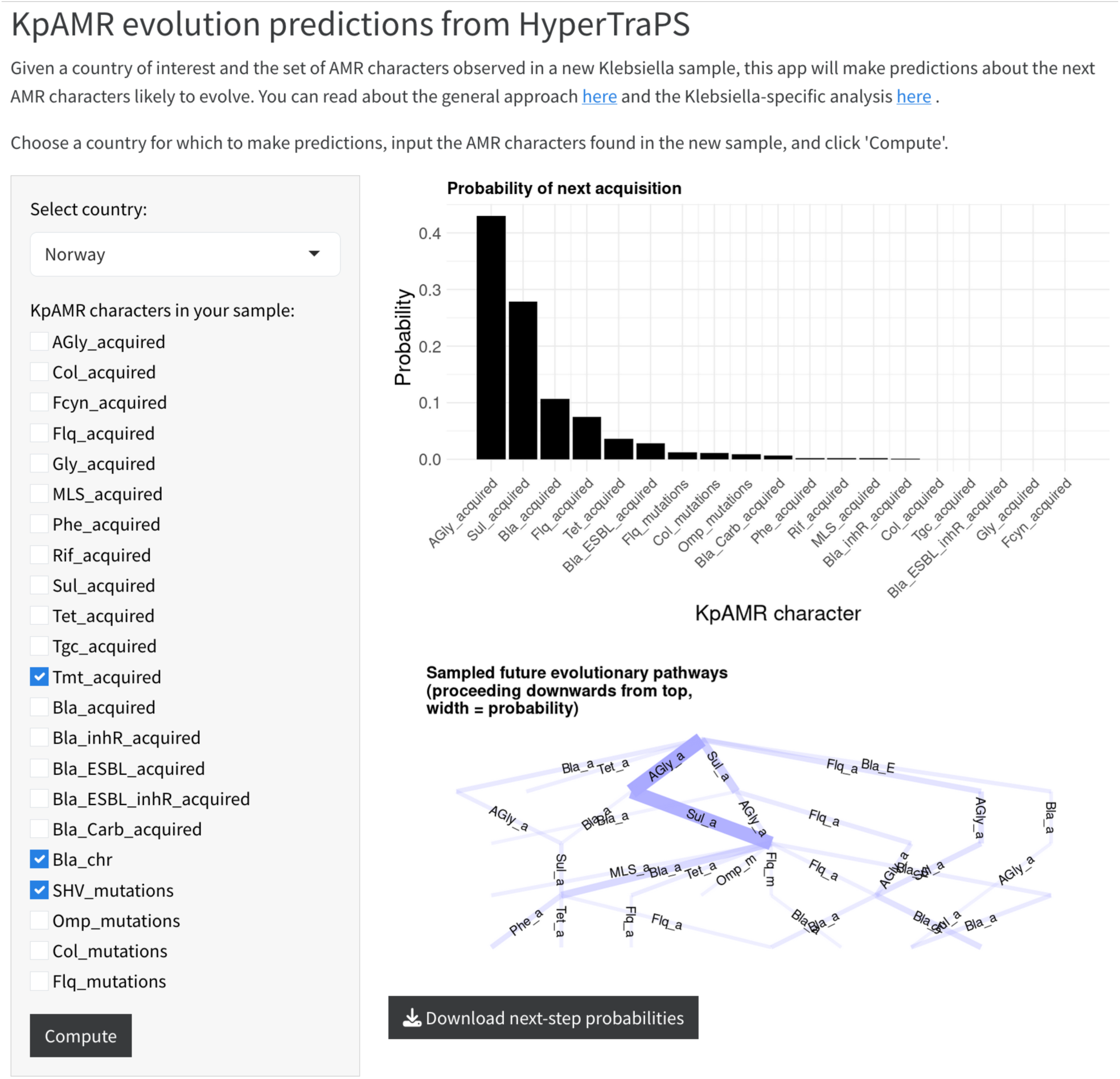
Shiny app for predictions of future KpAMR character acquisitions. A country of interest is specified, and the KpAMR profile of a sample given by specifying presence or absence of each of our characters. The graphical outputs then give the inferred probabilities of each next possible character acquisition (which can be downloaded in CSV format), as well as a transition graph describing the likely subsequent pathways for up to five acquisitions in the future. Available at https://stochasticbiology.shinyapps.io/amr-predict/.

## References

Aga, O. N. L., Brun, M., Giannakis, K., Dauda, K. A., Diaz-Uriarte, R., & Johnston, I. G. (2024). HyperTraPS-CT: Inference and prediction for accumulation pathways with flexible data and model structures. PLOS Computational Biology, 20(9), e1012393.

Aguilar-Bultet, L., García-Martín, A. B., Vock, I., Maurer Pekerman, L., Stadler, R., Schindler, R., Battegay, M., Stadler, T., Gómez-Sanz, E., & Tschudin-Sutter, S. (2023). Within-host genetic diversity of extended-spectrum beta-lactamase-producing Enterobacterales in long-term colonized patients. Nature Communications, 14, 8495. 10.1038/s41467-023-44285-w

Ahlmann-Eltze, C. (2025). ggupset: Combination Matrix Axis for ‘ggplot2’ to Create ‘UpSet’ Plots [R package].

Arel-Bundock, V., Enevoldsen, N., & Yetman, C. (2018). countrycode: An R package to convert country names and country codes. Journal of Open Source Software, 3(28), 848. 10.21105/joss.00848

Argimón, S., David, S., Underwood, A., Abrudan, M., Wheeler, N. E., Kekre, M., Abudahab, K., Yeats, C. A., Goater, R., Taylor, B., Harste, H., Muddyman, D., Feil, E. J., Brisse, S., Holt, K., Donado-Godoy, P., Ravikumar, K. L., Okeke, I. N., Carlos, C., … NIHR Global Health Research Unit on Genomic Surveillance of Antimicrobial Resistance. (2021). Rapid Genomic Characterization and Global Surveillance of Klebsiella Using Pathogenwatch. Clinical Infectious Diseases, 73(Supplement_4), S325–S335. 10.1093/cid/ciab784

Bates, D., Mächler, M., Bolker, B., & Walker, S. (2015). Fitting Linear Mixed-Effects Models Using lme4. Journal of Statistical Software, C7, 1–48. 10.18637/jss.v067.i01

Beerenwinkel, N., Däumer, M., Sing, T., Rahnenführer, J., Lengauer, T., Selbig, J., Hoffmann, D., & Kaiser, R. (2005). Estimating HIV evolutionary pathways and the genetic barrier to drug resistance. The Journal of Infectious Diseases, 1S1(11), 1953–1960.

Blanquart, F. (2019). Evolutionary epidemiology models to predict the dynamics of antibiotic resistance. Evolutionary Applications, 12(3), 365–383. 10.1111/eva.12753

Blomberg, B., Manji, K. P., Urassa, W. K., Tamim, B. S., Mwakagile, D. S. M., Jureen, R., Msangi, V., Tellevik, M. G., Holberg-Petersen, M., Harthug, S., Maselle, S. Y., & Langeland, N. (2007). Antimicrobial resistance predicts death in Tanzanian children with bloodstream infections: A prospective cohort study. BMC Infectious Diseases, 7, 43. 10.1186/1471-2334-7-43

Boyko, J. D., & Beaulieu, J. M. (2023). Reducing the biases in false correlations between discrete characters. Systematic Biology, 72(2), 476–488.

Casali, N., Nikolayevskyy, V., Balabanova, Y., Harris, S. R., Ignatyeva, O., Kontsevaya, I., Corander, J., Bryant, J., Parkhill, J., Nejentsev, S., Horstmann, R. D., Brown, T., C Drobniewski, F. (2014). Evolution and transmission of drug-resistant tuberculosis in a Russian population. Nature Genetics, 4C(3), Article 3. 10.1038/ng.2878

Clarke, E., Sherrill-Mix, S., & Dawson, C. (2016). ggbeeswarm: Categorical scatter (violin point) plots. CRAN: Contributed Packages. https://cir.nii.ac.jp/crid/1360019690351052672

Dauda, K. A., Aga, O. N. L., & Johnston, I. G. (2025). Clustering Large-Scale Biomedical Data to Model Dynamic Accumulation Processes in Disease Progression and Anti-Microbial Resistance Evolution. IEEE Access, 13, 13816–13831. 10.1109/ACCESS.2025.3527715

Dewar, A. E., Belcher, L. J., & West, S. A. (2025). A phylogenetic approach to comparative genomics. Nature Reviews Genetics, 2C(6), 395–405. 10.1038/s41576-024-00803-0

Diaz-Uriarte, R., & Herrera-Nieto, P. (2022). EvAM-Tools: Tools for evolutionary accumulation and cancer progression models. Bioinformatics, 38(24), 5457–5459.

Diaz-Uriarte, R., & Johnston, I. G. (2024). A picture guide to cancer progression and monotonic accumulation models: Evolutionary assumptions, plausible interpretations, and alternative uses (arXiv:2312.06824). arXiv. 10.48550/arXiv.2312.06824

Diaz-Uriarte, R., & Johnston, I. G. (2025). A Picture Guide to Cancer Progression and Evolutionary Accumulation Models: Systematic Critique, Plausible Interpretations, and Alternative Uses. IEEE Access, 13, 62306–62340. 10.1109/ACCESS.2025.3558392

Global Antimicrobial Resistance and Use Surveillance System (GLASS) Report 2022 (1st ed). (2022). World Health Organization.

Greenbury, S. F., Barahona, M., & Johnston, I. G. (2020). HyperTraPS: Inferring probabilistic patterns of trait acquisition in evolutionary and disease progression pathways. Cell Systems, 10(1), 39–51.

Gupta, S. K., Padmanabhan, B. R., Diene, S. M., Lopez-Rojas, R., Kempf, M., Landraud, L., & Rolain, J.-M. (2014). ARG-ANNOT, a new bioinformatic tool to discover antibiotic resistance genes in bacterial genomes. Antimicrobial Agents and Chemotherapy, 58(1), 212–220. 10.1128/AAC.01310-13

Han, J. H., Lapp, Z., Bushman, F., Lautenbach, E., Goldstein, E. J. C., Mattei, L., Hofstaedter, C. E., Kim, D., Nachamkin, I., Garrigan, C., Jain, T., Bilker, W., Wolford, H. M., Slayton, R. B., Wise, J., Tolomeo, P., & Snitkin, E. S. (2019). Whole-Genome Sequencing To Identify Drivers of Carbapenem-Resistant Klebsiella pneumoniae Transmission within and between Regional Long-Term Acute-Care Hospitals. Antimicrobial Agents and Chemotherapy, C3(11), 10.1128/aac.01622-19. 10.1128/aac.01622-19

Harris, C. R., Millman, K. J., van der Walt, S. J., Gommers, R., Virtanen, P., Cournapeau, D., Wieser, E., Taylor, J., Berg, S., Smith, N. J., Kern, R., Picus, M., Hoyer, S., van Kerkwijk, M. H., Brett, M., Haldane, A., del Río, J. F., Wiebe, M., Peterson, P., … Oliphant, T. E. (2020). Array programming with NumPy. Nature, 585(7825), 357–362. 10.1038/s41586-020-2649-2

Hennart, M., Guglielmini, J., Bridel, S., Maiden, M. C. J., Jolley, K. A., Criscuolo, A., & Brisse, S. (2022). A Dual Barcoding Approach to Bacterial Strain Nomenclature: Genomic Taxonomy of Klebsiella pneumoniae Strains. Molecular Biology and Evolution, 3S(7), msac135. 10.1093/molbev/msac135

Herencias, C., Álvaro-Llorente, L., Ramiro-Martínez, P., Fernández-Calvet, A., Muñoz-Cazalla, A., DelaFuente, J., Graf, F. E., Jaraba-Soto, L., Castillo-Polo, J. A., Cantón, R., San Millán, Á., & Rodríguez-Beltrán, J. (2024). β-lactamase expression induces collateral sensitivity in Escherichia coli. Nature Communications, 15(1), 4731. 10.1038/s41467-024-49122-2

Hetland, M. A. K., Winkler, M. A., Kaspersen, H. P., Håkonsholm, F., Bakksjø, R.-J., Bernhoff, E., Delgado-Blas, J. F., Brisse, S., Correia, A., Fostervold, A., Lam, M. M. C., Lunestad, B.-T., Marathe, N. P., Raffelsberger, N., Samuelsen, Ø., Sunde, M., Sundsfjord, A., Urdahl, A. M., Wick, R. R., … Holt, K. E. (2025). A genome-wide One Health study of Klebsiella pneumoniae in Norway reveals overlapping populations but few recent transmission events across reservoirs. Genome Medicine, 17(1), 42. 10.1186/s13073-025-01466-0

Hipólito, A., García-Pastor, L., Vergara, E., Jové, T., & Escudero, J. A. (2023). Profile and resistance levels of 136 integron resistance genes. Npj Antimicrobials and Resistance, 1(1), 13. 10.1038/s44259-023-00014-3

Holt, K. E., Wertheim, H., Zadoks, R. N., Baker, S., Whitehouse, C. A., Dance, D., Jenney, A., Connor, T. R., Hsu, L. Y., Severin, J., Brisse, S., Cao, H., Wilksch, J., Gorrie, C., Schultz, M. B., Edwards, D. J., Nguyen, K. V., Nguyen, T. V., Dao, T. T., … Thomson, N. R. (2015). Genomic analysis of diversity, population structure, virulence, and antimicrobial resistance in Klebsiella pneumoniae, an urgent threat to public health. Proceedings of the National Academy of Sciences, 112(27), E3574–E3581. 10.1073/pnas.1501049112

Johnston, I. G. (2026). Extracting useful information about reversible evolutionary processes from irreversible evolutionary accumulation models (arXiv:2601.13010). arXiv. 10.48550/arXiv.2601.13010

Johnston, I. G., & Diaz-Uriarte, R. (2024). A hypercubic Mk model framework for capturing reversibility in disease, cancer, and evolutionary accumulation modelling (p. 2024.06.27.600959). bioRxiv. 10.1101/2024.06.27.600959

Johnston, I. G., & Williams, B. P. (2016). Evolutionary Inference across Eukaryotes Identifies Specific Pressures Favoring Mitochondrial Gene Retention. Cell Systems, 2(2), 101–111. 10.1016/j.cels.2016.01.013

Kassambara, A. (2020). Ggpubr:“ggplot2” based publication ready plots. R Package Version 0.4. 0, 438.

Krieger, M. S., Denison, C. E., Anderson, T. L., Nowak, M. A., & Hill, A. L. (2020). Population structure across scales facilitates coexistence and spatial heterogeneity of antibiotic-resistant infections. PLOS Computational Biology, 1C(7), e1008010. 10.1371/journal.pcbi.1008010

Kurtz, S., Phillippy, A., Delcher, A. L., Smoot, M., Shumway, M., Antonescu, C., & Salzberg, S. L. (2004). Versatile and open software for comparing large genomes. Genome Biology, 5(2), R12. 10.1186/gb-2004-5-2-r12

Lam, M. M. C., Wick, R. R., Watts, S. C., Cerdeira, L. T., Wyres, K. L., & Holt, K. E. (2021). A genomic surveillance framework and genotyping tool for Klebsiella pneumoniae and its related species complex. Nature Communications, 12(1), 4188. 10.1038/s41467-021-24448-3

Lehtinen, S., Blanquart, F., Lipsitch, M., Fraser, C., & Collaboration, with the M. P. (2019). On the evolutionary ecology of multidrug resistance in bacteria. PLOS Pathogens, 15(5), e1007763. 10.1371/journal.ppat.1007763

Liu, P., Li, P., Jiang, X., Bi, D., Xie, Y., Tai, C., Deng, Z., Rajakumar, K., & Ou, H.-Y. (2012). Complete genome sequence of Klebsiella pneumoniae subsp. Pneumoniae HS11286, a multidrug-resistant strain isolated from human sputum. Journal of Bacteriology, 1S4(7), 1841–1842. 10.1128/JB.00043-12

Luo, X. G., Kuipers, J., C Beerenwinkel, N. (2023). Joint inference of exclusivity patterns and recurrent trajectories from tumor mutation trees. Nature Communications, 14(1), Article 1. 10.1038/s41467-023-39400-w

Maddison, W. P., & FitzJohn, R. G. (2015). The Unsolved Challenge to Phylogenetic Correlation Tests for Categorical Characters. Systematic Biology, C4(1), 127–136. 10.1093/sysbio/syu070

Mapunjo, S., Mbwasi, R., Nkiligi, E. A., Wilbroad, A., Francis, E. N., Msovela, K., Yahya, T., Mpembeni, R., Masunga, E., Nkungu, K., Saitoti, S., Lusaya, E., & Konduri, N. (2025). National consumption of antimicrobials in Tanzania: 2020–2022. JAC-Antimicrobial Resistance, 7(2), dlaf026. 10.1093/jacamr/dlaf026

Marakeby, H., Badr, E., Torkey, H., Song, Y., Leman, S., Monteil, C. L., Heath, L. S., & Vinatzer, B. A. (2014). A System to Automatically Classify and Name Any Individual Genome-Sequenced Organism Independently of Current Biological Classification and Nomenclature. PLOS ONE, S(2), e89142. 10.1371/journal.pone.0089142

Moen, M. T., & Johnston, I. G. (2023). HyperHMM: Efficient inference of evolutionary and progressive dynamics on hypercubic transition graphs. Bioinformatics, 3S(1), btac803.

Moyo, S. J., Manyahi, J., Blomberg, B., Tellevik, M. G., Masoud, N. S., Aboud, S., Manji, K., Roberts, A. P., Hanevik, K., Mørch, K., & Langeland, N. (2020). Bacteraemia, Malaria, and Case Fatality Among Children Hospitalized With Fever in Dar es Salaam, Tanzania. Frontiers in Microbiology, 11, 2118. 10.3389/fmicb.2020.02118

Munk, P., Brinch, C., Møller, F. D., Petersen, T. N., Hendriksen, R. S., Seyfarth, A. M., Kjeldgaard, J. S., Svendsen, C. A., van Bunnik, B., Berglund, F., Larsson, D. G. J., Koopmans, M., Woolhouse, M., & Aarestrup, F. M. (2022). Genomic analysis of sewage from 101 countries reveals global landscape of antimicrobial resistance. Nature Communications, 13(1), 7251. 10.1038/s41467-022-34312-7

Murray, C. J. L., Ikuta, K. S., Sharara, F., Swetschinski, L., Aguilar, G. R., Gray, A., Han, C., Bisignano, C., Rao, P., Wool, E., Johnson, S. C., Browne, A. J., Chipeta, M. G., Fell, F., Hackett, S., Haines-Woodhouse, G., Hamadani, B. H. K., Kumaran, E. A. P., McManigal, B., … Naghavi, M. (2022). Global burden of bacterial antimicrobial resistance in 2019: A systematic analysis. The Lancet, 3SS(10325), 629–655. 10.1016/S0140-6736(21)02724-0

Naghavi, M., Vollset, S. E., Ikuta, K. S., Swetschinski, L. R., Gray, A. P., Wool, E. E., Aguilar, G. R., Mestrovic, T., Smith, G., Han, C., Hsu, R. L., Chalek, J., Araki, D. T., Chung, E., Raggi, C., Hayoon, A. G., Weaver, N. D., Lindstedt, P. A., Smith, A. E., … Murray, C. J. L. (2024). Global burden of bacterial antimicrobial resistance 1990–2021: A systematic analysis with forecasts to 2050. The Lancet, 404(10459), 1199–1226. 10.1016/S0140-6736(24)01867-1

Olesen, S. W., Barnett, M. L., MacFadden, D. R., Brownstein, J. S., Hernández-Díaz, S., Lipsitch, M., & Grad, Y. H. (2018). The distribution of antibiotic use and its association with antibiotic resistance. eLife, 7, e39435. 10.7554/eLife.39435

Onken, A., Moyo, S., Miraji, M. K., Bohlin, J., Marijani, M., Manyahi, J., Kibwana, K. O., Müller, F., Jenum, P. A., Abeid, K. A., Reimers, M., Langeland, N., Mørch, K., & Blomberg, B. (2024). Predominance of multidrug-resistant Salmonella Typhi genotype 4.3.1 with low-level ciprofloxacin resistance in Zanzibar. PLOS Neglected Tropical Diseases, 18(4), e0012132. 10.1371/journal.pntd.0012132

Onken, A., Said, A. K., Jørstad, M., Jenum, P. A., & Blomberg, B. (2015). Prevalence and Antimicrobial Resistance of Microbes Causing Bloodstream Infections in Unguja, Zanzibar. PLoS ONE, 10(12), e0145632. 10.1371/journal.pone.0145632

Pál, C., Papp, B., & Lázár, V. (2015). Collateral sensitivity of antibiotic-resistant microbes. Trends in Microbiology, 23(7), 401–407. 10.1016/j.tim.2015.02.009

R Core Team. (2022). R: A language and environment for statistical computing. R Foundation for Statistical Computing, Vienna, Austria. 2012.

Renz, J., Dauda, K. A., Aga, O. N. L., Diaz-Uriarte, R., Löhr, I. H., Blomberg, B., & Johnston, I. G. (2024). Evolutionary accumulation modelling in AMR: Machine learning to infer and predict evolutionary dynamics of multi-drug resistance (arXiv:2411.00219). arXiv. 10.48550/arXiv.2411.00219

Renz, J., Dauda, K. A., Aga, O. N. L., Diaz-Uriarte, R., Löhr, I. H., Blomberg, B., & Johnston, I. G. (2025). Evolutionary accumulation modeling in AMR: Machine learning to infer and predict evolutionary dynamics of multi-drug resistance. mBio, 1C(6), e00488–25. 10.1128/mbio.00488-25

Revell, L. J. (2012). phytools: An R package for phylogenetic comparative biology (and other things). Methods in Ecology and Evolution, (2), 217–223.

Rudd, K. E., Johnson, S. C., Agesa, K. M., Shackelford, K. A., Tsoi, D., Kievlan, D. R., Colombara, D. V., Ikuta, K. S., Kissoon, N., Finfer, S., Fleischmann-Struzek, C., Machado, F. R., Reinhart, K. K., Rowan, K., Seymour, C. W., Watson, R. S., West, T. E., Marinho, F., Hay, S. I., … Naghavi, M. (2020). Global, regional, and national sepsis incidence and mortality, 1990–2017: Analysis for the Global Burden of Disease Study. The Lancet, 3S5(10219), 200–211. 10.1016/S0140-6736(19)32989-7

Saitou, N., & Nei, M. (1987). The neighbor-joining method: A new method for reconstructing phylogenetic trees. Molecular Biology and Evolution, 4(4), 406–425.

Sangeda, R. Z., Saburi, H. A., Masatu, F. C., Aiko, B. G., Mboya, E. A., Mkumbwa, S., Bitegeko, A., Mwalwisi, Y. H., Nkiligi, E. A., Chambuso, M., Sillo, H. B., Fimbo, A. M., & Horumpende, P. G. (2021). National Antibiotics Utilization Trends for Human Use in Tanzania from 2010 to 2016 Inferred from Tanzania Medicines and Medical Devices Authority Importation Data. Antibiotics, 10(10), 1249. 10.3390/antibiotics10101249

Schill, R., Klever, M., Rupp, K., Hu, Y. L., Lösch, A., Georg, P., Pfahler, S., Vocht, S., Hansch, S., Wettig, T., Grasedyck, L., & Spang, R. (2024). Reconstructing Disease Histories in Huge Discrete State Spaces. KI - Künstliche Intelligenz. 10.1007/s13218-023-00822-9

Schill, R., Solbrig, S., Wettig, T., & Spang, R. (2020). Modelling cancer progression using mutual hazard networks. Bioinformatics, 3C(1), 241–249.

Schliep, K. P. (2011). phangorn: Phylogenetic analysis in R. Bioinformatics, 27(4), 592–593. 10.1093/bioinformatics/btq706

Sköld, O. (2001). Resistance to trimethoprim and sulfonamides. Veterinary Research, 32(3– 4), 261–273. 10.1051/vetres:2001123

Slowikowski, K. (2021). ggrepel: Automatically Position Non-Overlapping Text Labels with’ggplot2’. R package version 0.S. 1, 2021.

Sokal, R. R., & Michener, C. D. (1958). A statistical method for evaluating systematic relationships. University of Kansas Science Bulletin, 38, 1409.

Szybalski, W., & Bryson, V. (1952). Genetic studies on microbial cross resistance to toxic agents i. Journal of Bacteriology, C4(4), 489–499. 10.1128/jb.64.4.489-499.1952

Tsang, K. K., Lam, M. M. C., Wick, R. R., Wyres, K. L., Bachman, M., Baker, S., Barry, K., Brisse, S., Campino, S., Chiaverini, A., Cirillo, D. M., Clark, T., Corander, J., Corbella, M., Cornacchia, A., Cuénod, A., D’Alterio, N., Di Marco, F., Donado-Godoy, P., … KlebNET-GSP AMR Genotype-Phenotype Group. (2024). Diversity, functional classification and genotyping of SHV β-lactamases in Klebsiella pneumoniae. Microbial Genomics, 10(10), 001294. 10.1099/mgen.0.001294

Wickham, H. (2016). ggplot2: Elegant Graphics for Data Analysis. Springer-Verlag. https://link.springer.com/book/10.1007/978-0-387-98141-3

Wickham, H., Averick, M., Bryan, J., Chang, W., McGowan, L., François, R., Grolemund, G., Hayes, A., Henry, L., Hester, J., Kuhn, M., Pedersen, T., Miller, E., Bache, S., Müller, K., Ooms, J., Robinson, D., Seidel, D., Spinu, V., … Yutani, H. (2019). Welcome to the Tidyverse. Journal of Open Source Software, 4(43), 1686. 10.21105/joss.01686

Wickham, H., François, R., Henry, L., Müller, K., & Vaughan, D. (2023). Dplyr: A grammar of data manipulation. R package version 1.1. 2. Computer Software.

Williams, B. P., Johnston, I. G., Covshoff, S., & Hibberd, J. M. (2013). Phenotypic landscape inference reveals multiple evolutionary paths to C4 photosynthesis. eLife, 2, e00961. 10.7554/eLife.00961

Wu, Y., Jiang, T., He, X., Shao, J., Wu, C., Mao, W., Jia, H., He, F., Kong, Y., Wu, J., Sun, Q., Sun, L., Draz, M. S., Xie, X., Zhang, J., & Ruan, Z. (2023). Global Phylogeography and Genomic Epidemiology of Carbapenem-Resistant blaOXA-232–Carrying Klebsiella pneumoniae Sequence Type 15 Lineage. Emerging Infectious Diseases, 2S(11), 2246–2256. 10.3201/eid2911.230463

Wyres, K. L., & Holt, K. E. (2018). Klebsiella pneumoniae as a key trafficker of drug resistance genes from environmental to clinically important bacteria. Current Opinion in Microbiology, 45, 131–139. 10.1016/j.mib.2018.04.004

Wyres, K. L., Lam, M. M. C., & Holt, K. E. (2020). Population genomics of Klebsiella pneumoniae. Nature Reviews. Microbiology, 18(6), 344–359. 10.1038/s41579-019-0315-1

